# Guiding oligodendrocyte precursor cell maturation with urokinase plasminogen activator-degradable elastin-like protein hydrogels

**DOI:** 10.1101/2020.07.28.224899

**Authors:** Edi Meco, W. Sharon Zheng, Anahita H. Sharma, Kyle J. Lampe

## Abstract

Demyelinating injuries and diseases, like multiple sclerosis, affect millions of people worldwide. Oligodendrocyte precursor cells (OPCs) have the potential to repair demyelinated tissue because they can both self-renew and differentiate into oligodendrocytes (OLs), the myelin producing cells of the central nervous system (CNS). Cell-matrix interactions impact OPC differentiation into OLs, but the process is not fully understood. Biomaterial hydrogel systems help to elucidate cell-matrix interactions because they can mimic specific properties of native CNS tissue in an *in vitro* setting. We investigated whether OPC maturation into OLs is influenced by interacting with a urokinase plasminogen activator (uPA) degradable extracellular matrix (ECM). uPA is a proteolytic enzyme that is transiently upregulated in the developing rat brain, with peak uPA expression correlating with an increase in myelin production *in vivo*. OPC-like cells isolated through the Mosaic Analysis with Double Marker technique (MADM OPCs) produced low molecular weight uPA in culture. MADM OPCs were encapsulated into two otherwise similar elastin-like protein (ELP) hydrogel systems: one that was uPA degradable and one that was non-degradable. Encapsulated MADM OPCs had similar viability, proliferation, and metabolic activity in uPA degradable and non-degradable ELP hydrogels. Expression of OPC maturation-associated genes, however, indicated that uPA degradable ELP hydrogels promoted MADM OPC maturation although not sufficiently for these cells to differentiate into OLs.

**Graphical Abstract – For table of contents only:** 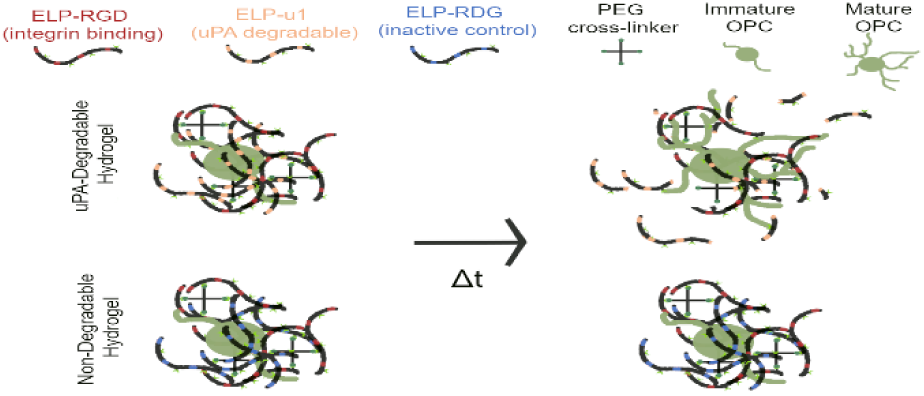

## 1 Introduction

Biomaterial hydrogel systems are effective cell carriers that support high transplanted cell viability in the context of cytotoxic extracellular environments^1,2^. In order to improve the disease treatment potential of hydrogel systems, current research efforts focus on designing biomaterial systems that interact with encapsulated cells to induce desired behavior^3^. Demyelinating injuries and diseases, like multiple sclerosis (MS), affect millions of people worldwide^4^. The progression of demyelinating injuries and diseases damages central nervous system (CNS) oligodendrocytes, which extend processes to wrap an insulating myelin sheath around neuronal axons. This myelin supports nerves in the CNS by helping to propagate synaptic signals and provide trophic support to ensheathed axons^5,6^. Oligodendrocyte precursor cells (OPCs) have the potential to repair damaged tissue because they can both self-renew and differentiate into OLs. However, endogenous OPCs are minimally able to repair demyelinated tissue because, in part, the extracellular matrix (ECM) within the these lesions is not conducive for OPCs to migrate, differentiate, and replace damaged OLs^7,8^. A key factor in the development of treatments for demyelinated tissue is to design hydrogel systems that promote OPC maturation into OLs.

OPC maturation into OLs is affected by many extrinsic factors, such as ECM stiffness and the presence of fibronectin^8–10^. In this study we investigated the influence of urokinase plasminogen activator (uPA) enzymatic activity and 3D uPA-responsive hydrogels on OPC maturation. uPA is a serine protease enzyme involved in ECM remodeling, and acts as an activator of the plasminogen system^11^. uPA expression *in vivo* indicates that it plays a key role in the OPC maturation process. During rat development, uPA is transiently expressed in myelinating regions (corpus collosum and fimbria) of the brain^12^. Peak uPA expression in these regions occurs around postnatal day 14 (P14), and it is lost by adulthood (P56)^12^. Peak myelin development (P10-17) in many myelinated regions of the rat brain coincides with peak uPA expression^13^. Further evidence that indicates uPA may play a role in OPC maturation comes from the impact it has on cell morphology. OPC maturation into OLs is marked by a morphological increase in the number of processes extended and branching of those processes^14^. Although no study has tested the effects of uPA expression changes on OPC morphology, its expression affects cellular process extension. Pharmacological inhibition of uPA in dorsal root ganglia explants encapsulated within a Matrigel matrix reduced the average axon length by 66 %^15^.

uPA may also impact OPC maturation indirectly through its role as an activator of the plasminogen system, which upregulates the expression of many matrix metalloproteinases (MMPs)^11^. In particular, MMP-9 may promote OPC maturation because it is upregulated in the corpus collosum and optic nerve regions during developmental myelination in mice^16,17^. MMP-9 also plays a significant role in the subsequent remyelination of demyelinated lesions post injury. MMP-9 null mice were found to have reduced remyelination 1-week post lysolecithin-induced demyelination spinal cord injury when compared to wild type mice^16^. The reduction in remyelination correlated with less mature OLs present within the injury lesion of MMP-9 null mice compared to wild type mice^16^. Taken together, previous findings indicate that uPA may affect OPC maturation either directly through ECM remodeling and/or indirectly through the plasminogen activation of MMP-9. Hydrogel systems that promote uPA expression could therefore also stimulate OPC maturation.

Hydrogels made from cross-linked elastin-like proteins (ELPs) are suitable for cell encapsulation^18–22^. Engineered ELPs are designed based on native tropoelastin, and consist of repeating the penta-peptide sequence valine-proline-glycine-x-glycine (VPGxG), where x is any amino acid guest residue except for proline. ELPs exhibit a unique lower critical solution temperature (LCST) transition^23–25^. Above a sequence dependent transition temperature (T_t_) the protein separates into a protein-rich coacervate phase. Bioactivity, adjunct to the elastin component of the penta-peptide sequence, is incorporated into ELPs by including other peptide sequences in the protein^19,22,26^. In this study, we use three protein sequences composed of the same elastin-like penta-peptide repeating sequence and differing bioactive peptide sequences: ELP-u1 contains a uPA enzymatically cleavable sequence, ELP-RGD contains a sequence for integrin binding and ELP-RDG contains a sequence that is a scramble of RGD with no integrin-binding activity (Figure 1)^26^. The similarity in the amino acid content (> 98 %) between these three ELPs allows for the creation of hydrogels with adjustable bioactivity (for instance, integrin binding and degradation) independent from the resulting hydrogel mechanical properties^26,27^. Hydrogel bioactivity can be tuned by adjusting the ratios of each ELP sequence used, and mechanical properties can be tuned by adjusting the total protein concentration and cross-linking ratio. OPC maturation is impacted by both the macromolecular bioactivity and the mechanical properties of the surrounding ECM^8– 10,28,29^. Independent control of ELP hydrogel bioactivity and mechanical properties makes it a suitable *in vitro* model to investigate the impact of extrinsic factors on OPC behavior.

**Figure 1:**
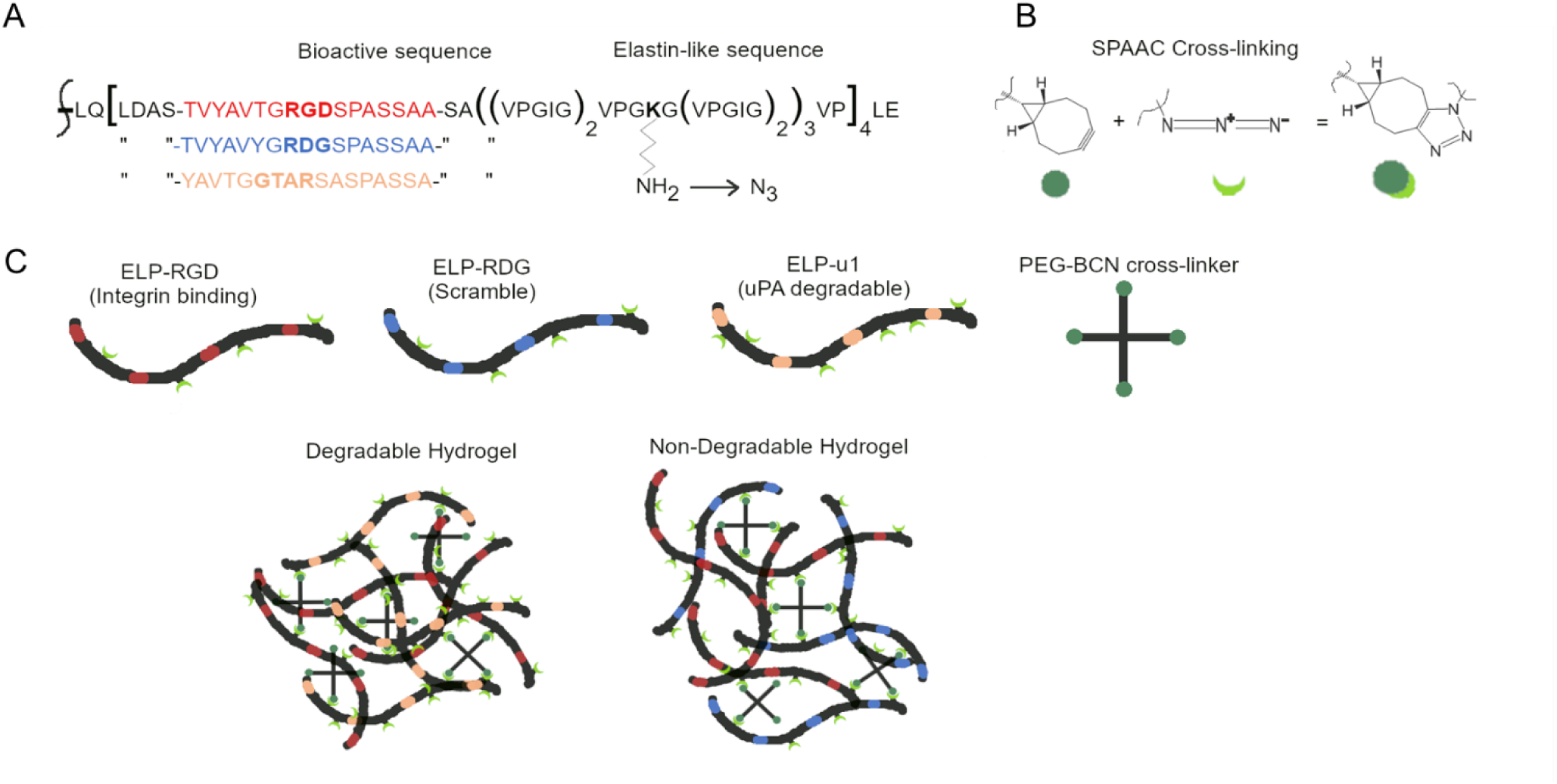
Formation of urokinase plasminogen activator (uPA) degradable and non-degradable elastin-like protein (ELP) hydrogels using strain-promoted azide alkyne cycloaddition (SPAAC) chemistry. (A) Three ELP sequences with different bioactivities were used: ELP-RGD promotes integrin binding, ELP-RDG is a non-bioactive scramble, and ELP-u1 is uPA cleavable. Lysine residues on ELP were converted into azide groups. (B) ELPs were cross-linked with 4 arm-PEG-BCN using SPAAC chemistry. (C) uPA degradable hydrogels consisted of a 50/50 % mix of ELP-RGD and ELP-u1, while non-degradable hydrogels consisted of a 50/50 % mix of ELP-RGD and ELP-RDG. Both hydrogels contain identical concentrations of integrin-binding RGD motifs.

The goal of this study was to determine if OPC maturation could be influenced by incorporating uPA enzymatic degradability into the ELP hydrogel network. Immortalized OPC-like cells isolated from mice using the mosaic analysis with double markers method (MADM OPC) were encapsulated in both uPA degradable and non-degradable ELP hydrogels^30,31^. MADM OPCs are NF1 and p53 gene knockouts that have delayed maturation *in vivo*. No current *in vitro* method to differentiate them into mature OLs exists^30,31^. We aimed to test whether or not MADM OPCs expressed uPA, to determine if encapsulating MADM OPCs in uPA degradable hydrogels would impact MADM OPC uPA expression, and to determine if uPA degradable hydrogels influenced MADM OPC maturation state.

## 2 Experimental Methods

### 2.1 Materials

All materials and reagents were purchased from Fisher Scientific, VWR, or Sigma-Aldrich and used without further modification unless otherwise noted.

### 2.2 ELP expression and purification

Three previously established ELP sequences were used (Table 1)^26^: ELP expression in *Escherichia coli* strain BL21(DE3) was induced with β-isopropyl thiogalactoside and purified as previously described without modification^32^. Protein yields were 50-130 mg/L.

**Table 1:**
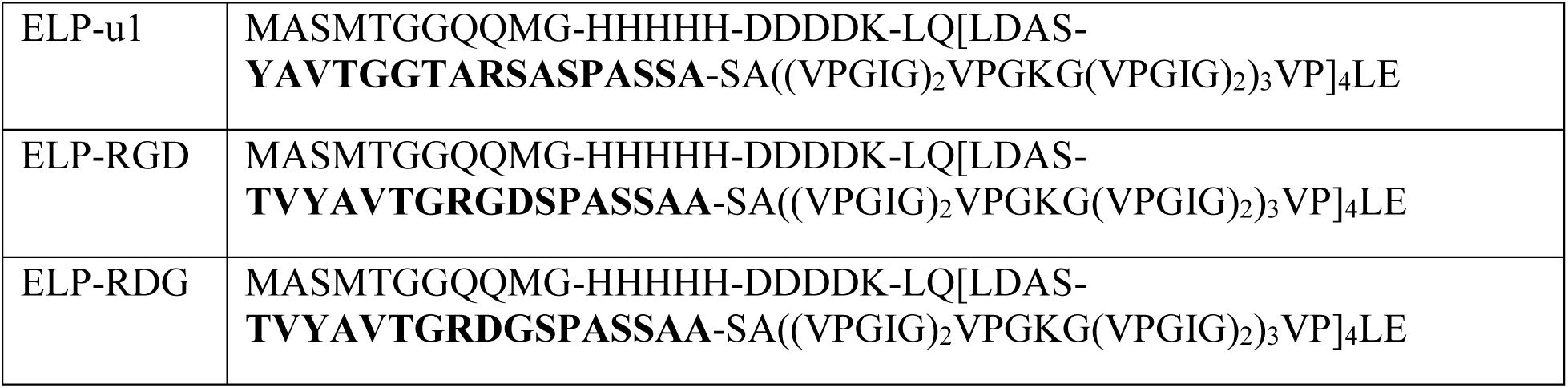
The amino acid sequences of ELP-u1, -RGD, and -RDG. Each ELP amino acid sequence contains a T7 tag, His Tag, enterokinase cleavage sequence, elastin-like sequence and a bioactive sequence (in bold).

|  |  |
| --- | --- |
| ELP-u1 | MASMTGGQQMG-HHHHH-DDDDK-LQ[LDAS-<br>**YAVTGGTARSASPASSA**-SA((VPGIG)<sub>2</sub>VPGKG(VPGIG)<sub>2</sub>)<sub>3</sub>VP]<sub>4</sub>LE |
| ELP-RGD | MASMTGGQQMG-HHHHH-DDDDK-LQ[LDAS-<br>**TVYAVTGRGDSPASSAA**-SA((VPGIG)<sub>2</sub>VPGKG(VPGIG)<sub>2</sub>)<sub>3</sub>VP]<sub>4</sub>LE |
| ELP-RDG | MASMTGGQQMG-HHHHH-DDDDK-LQ[LDAS-<br>**TVYAVTGRDGSPASSAA**-SA((VPGIG)<sub>2</sub>VPGKG(VPGIG)<sub>2</sub>)<sub>3</sub>VP]<sub>4</sub>LE |

### 2.3 ELP azide functionalization and characterization

Azide functionalization of primary amines on ELP was performed as previously described with minor modifications^33,34^. ELP (25 mg/ml) was dissolved in reaction buffer (potassium carbonate (10 mg/ml) in deionized (DI) water). The ELP solution was mixed with 100 μl copper (II) chloride solution (1 mg/ml in reaction buffer) in an argon purged round bottom flask (RBF). 1H-imidazole-1-sulfonyl azide HCl (25 mg/ml in reaction buffer, Enamine) was added to achieve a lysine residue amine to azide stoichiometric ratio of 14:4. The reaction was incubated for 24 hrs at 25°C with agitation. Azide-functionalized ELP was purified via dialysis using cellulose ester (CE) tubing (3.5-5 kDa molecular weight cut-off (MWCO)). Dialyzed ELP solution was frozen (−80 °C) and lyophilized to yield the final product, a white caked protein with slight red tint. Azide functionalization was confirmed with Fourier-transform infrared spectroscopy (FTIR, Supplemental Figure 1, F Perkin Elmer Frontier MIR/NIR spectrometer) and quantified with electrospray ionization-mass spectrometry (ESI-MS, Supplemental Figure 2, Thermo Scientific Q Exactive HF-X).

**Figure 2:**
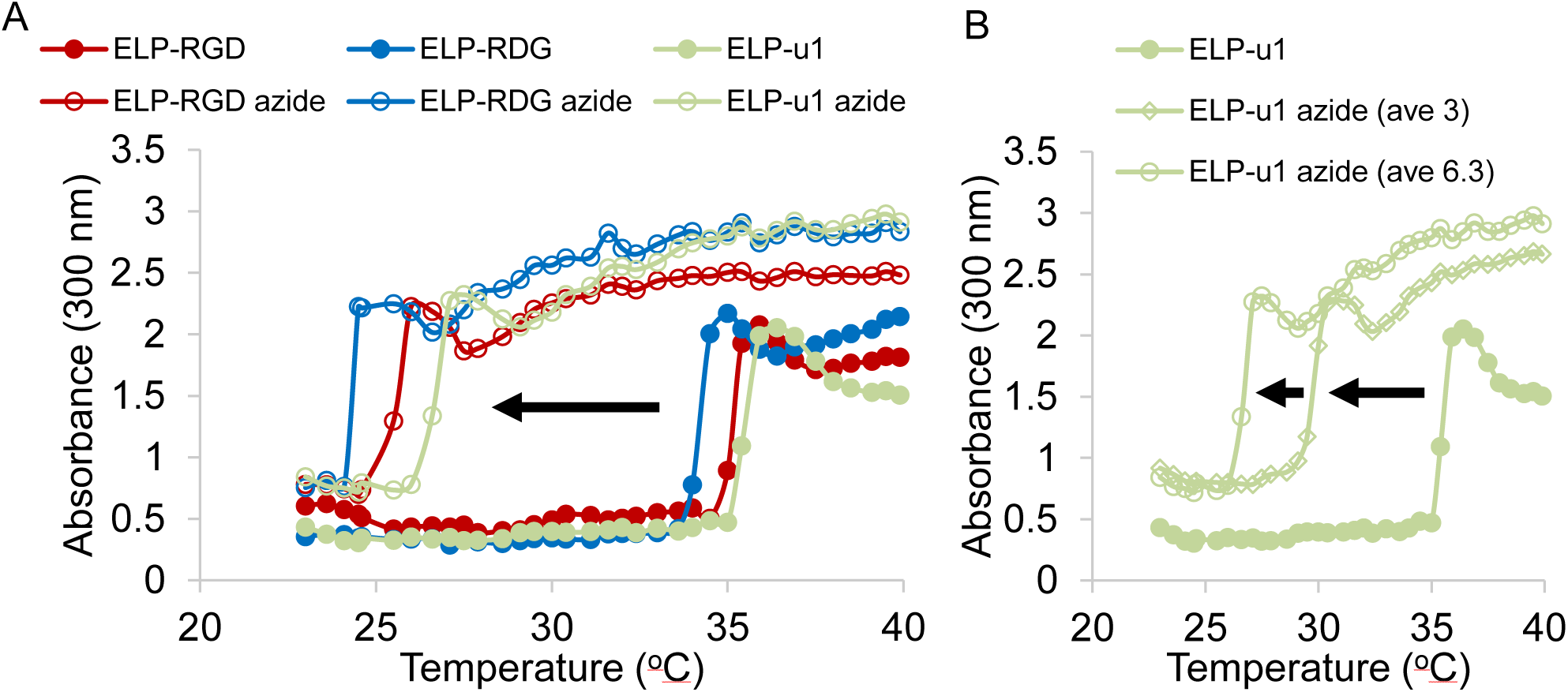
ELP LCST shift from azide reaction. (A) The LCST of all ELP sequences decreased when modified with azide groups. (B) The ELP LCST shift was dependent on the number of primary amines converted into azides.

### 2.4 PEG-BCN synthesis and characterization

4-arm poly(ethylene glycol) amine (PEG-amine) was functionalized with strained alkyne rings (BCN) as previously described with minor modifications^35^. PEG-amine (10 kDa, Laysan Bio) was dissolved in dimethylformamide (DMF) at a concentration of 200 mg/ml. In an argon purged RBF, PEG-amine was mixed with (1R, 8S, 9s)-Bicyclo[6.1.0]non-4-yn-9-ylmethyl N-succinimidyl carbonate (BCN-NHS) at a stoichiometric ratio of 1.5:1 BCN-NHS to amine groups. N,N-Diisopropylethylamine (DIEA) was added to the reaction mixture to achieve a 4:1 molar ratio DIEA to amine groups. The reaction mixture was stirred for 24 hrs at 25°C. The reaction mixture was diluted with DI water (20/80 v/v % reaction mixture/DI water), and the reacted product (PEG-BCN) was purified via dialysis using CE tubing (500-1,000 Da MWCO). Dialyzed PEG-BCN solution was frozen (−80 °C) and lyophilized to yield a white powder. PEG-BCN was dissolved in phosphate buffered saline (PBS, 200 mg/ml) and stored frozen (−80°C). PEG-BCN strained alkyne functionalization was quantified via proton nuclear magnetic resonance (HNMR, Varian INOVA 500 MHz Spectrometer) by comparing the PEG backbone peak (3.63 ppm) integral value to those of the attached BCN (0.92, 1.34, 1.57, and 2.24 ppm).

### 2.5 ELP LCST measurements

ELP and ELP azide were dissolved in PBS (50 mg/ml). 300 μl of each ELP solution and a PBS blank were pipetted into a 96-well plate. Absorbance was measured at 300 nm (BMG LABTECH CLARIOstar monochromatic microplate reader) for temperatures from 23 – 40 °C at 0.5 °C increments with a 0.5 °C/min temperature ramp. Samples were shaken at 300 rpm before each measurement. The T_t_ was calculated as the center point of the temperature range where the absorbance rapidly increased.

### 2.6 MADM OPC culture

MADM OPCs were cultured on T-75 plates, coated with polyornithine, using base media, which consisted of DMEM, high glucose (4.5 g/L), pyruvate supplemented with Penicillin-Streptomycin (1x), B27 supplement (1x), and N2 supplement (1x). At 90 % confluency MADM OPCs were extracted using trypsin (0.5x), counted using a Bright-Line Hemocytometer, and suspended in PBS. For monolayer cell cultures MADM OPCs were seeded at a density of 1×10^4^ cells/cm^2^, and hydrogel encapsulations were performed at a concentration of 5×10^5^ cells/ml. Media was changed every two days.

### 2.7 Hydrogel formation

Azide-functionalized ELP and PEG-BCN were dissolved in PBS and kept on ice prior to mixing. Gelation was performed by mixing the appropriate ELP and PEG-BCN solutions to form 10 wt % ELP hydrogels with a 0.5:1 BCN to azide stoichiometric cross-linking ratio; uPA degradable hydrogels consisted of a 50/50 % mixture of ELP-RGD and ELP-u1 sequences, and non-degradable gels consisted of a 50/50 % mixture of ELP-RGD and ELP-RDG sequences.

For MADM OPC encapsulations ELP was sterilized with ultraviolet light (UV) irradiation prior to dissolving in PBS. PEG-BCN solutions were sterilized by passing through a syringe filter (0.2 μm). Hydrogels were formed by mixing ELP, MADM OPC and PEG-BCN solutions together and pipetting 25 μl into 5 mm diameter molds (final concentrations as previously stated). Gelation proceeded by incubating solutions at room temperature for 30 min, followed by 30 min at 37 °C. Hydrogels were immersed in 700 μl base media.

### 2.8 Rheological measurements

Rheological measurements were conducted on an Anton Paar MCR 302 rheometer using a 25 mm cone and plate measuring probe with the stage set to 22°C. Azide-functionalized ELP and PEG-BCN solutions were kept on ice. Gelation was performed on the rheometer stage with the final concentrations as previously stated. 60 μl of the gel mixture was pipetted onto the stage for measurements. Time sweeps were conducted for 20 min with constant 1% strain and 1 Hz frequency. Strain sweeps were conducted with a logarithmic ramp from 0.01-1,000 % at a constant 1 Hz frequency. Frequency sweeps were conducted with a logarithmic ramp from 0.01-10 Hz at a constant 5 % strain. Stress relaxation was measured by holding the ELP hydrogels at 10 % strain for 15 min.

### 2.9 Zymography

The poly(acrylamide) gels were formed by immobilizing ELP onto the separating gel of a sodium dodecyl sulfate-polyacrylamide gel electrophoresis (SDS-PAGE) system. ELP was mixed with enough tetrakis(hydroxymethyl) phosphonium chloride (THPC) to achieve a 1:10 stoichiometric ratio of THPC hydroxyl groups to ELP primary amines, and added to the separating gel solution (2 mg/ml final ELP concentration). The stacking and separating gels were 3.9 and 7.5 % acrylamide, respectively.

Samples were collected as follows prior to use. Commercial uPA was diluted in PBS after purchase. MADM OPC base media was concentrated 30x using a protein concentrator PES (10,000 MWCO). Cultured MADM OPC media, collected from both monolayers and hydrogels, was centrifuged at 1,000 rpm for 5 min, decanted to remove any pelleted cell debris, and concentrated 30x. MADM OPC monolayer was collected by immersing a cell pellet in lysis buffer at a concentration of 1.1×10^8^ cells/ml. Encapsulated MADM OPCs were grown in ELP hydrogels for 4 days and collected by immersing in 700 μl lysis buffer. Hydrogels were homogenized with a pestle before use. 3 μl of each sample was mixed with 12 μl 5x non-reducing buffer (4 % SDS, 20 % glycerol, 0.01 % bromophenol blue, 125 mM TrisHCl (pH 6.8)). 15 μl of each sample were loaded into the acrylamide stacking gel wells, and electrophoresis was performed at 140 volts for 40 min.

Gels were washed twice for 30 min in washing buffer (2.5 % Triton X-100 in DI water)^36^. This was followed by a 10 min rinse at 37°C in incubation buffer (0.1 M Glycine pH 8.3 in DI water) with agitation. Gels were then incubated in incubation buffer for 24 hrs at 37°C. Gels were stained with Coomassie blue solution (30/10/60 % methanol/acetic acid/water, Coomassie blue concentration of 5 mg/ml) for 1 hr and destained (30/10/60 % methanol/acetic acid/water) prior to imaging.

### 2.10 MADM OPC degradation of ELP hydrogels

MADM OPCs were encapsulated in uPA degradable and non-degradable hydrogels as previously described. Hydrogels were imaged on days 1, 2 and 3 (Ziess Axio Observer fluorescent microscope) using green channel to visualize the green fluorescent protein positive (GFP+) OPCs. Tile images were stitched together and oriented in ImageJ software.

### 2.11 MADM OPC Imaging

MADM OPCs were encapsulated in uPA degradable and non-degradable hydrogels as previously described and cultured for up to 4 days. MADM OPC viability was measured on day 3 by incubating hydrogels with Dead stain (2 μM ethidium homodimer solution in PBSG) for 30 min at 37°C followed by immediate imaging. MADM OPC proliferation was measured on day 3 by staining hydrogels using the Click-iT EdU Imaging kit (Life Technologies) with minor modification. Briefly, cell-laden hydrogels were incubated in EdU solution for 1 hr at 37°C. They were then fixed in 3.7 % paraformaldehyde solution for 1 hr at 37°C. Two 5 min washes with 3 % bovine serum albumin (BSA) in PBS were conducted after fixation and in-between the following steps: Fixed hydrogels were permeabilized in 0.5 % Triton X-100 for 20 min, incubated in Click-iT reaction buffer for 30 min. MADM OPCs were stained for F-actin on day 4 with Alexa Fluor 568 phalloidin (Invitrogen) by incubating for 30 min in a phalloidin solution (2 U/ml in PBS with 1 % BSA) after fixing and permeabilizing as previously described. MADM OPC nuclei were stained with DAPI by incubating hydrogels in 300 nM DAPI solution for 30 min. Fixed hydrogels were stored in PBS at 4°C prior to imaging. All images were collected on a Leica SP8 X confocal microscope and processed with ImageJ software. Hydrogels were sealed between two coverslips to prevent dehydration. Z-stacks of each hydrogel sample were taken at 3 (Dead stain) and 5 (EdU stain) xy-locations. Max intensity projections were made of 15 (Dead stain) and 20 (EdU stain) μm sections along the z-axis, with 15 (Dead Stain) and 20 (EdU stain) μm gaps between sections. Viability was quantified by counting the number of GFP+ cells against Dead stained cells in each max projection. Proliferation was quantified by counting the number of EdU+ cells against DAPI and GFP+ cells in each max projection.

### 2.12 ATP and DNA quantification

MADM OPCs were encapsulated in ELP hydrogels as previously described and cultured for up to 5 days. Hydrogels were collected in 700 μl lysis buffer and homogenized with a pestle. ATP was measured using CellTiter-Glo luminescent assay (Promega) using manufacturer’s protocol. Homogenized solutions were sonicated for 10 seconds and DNA was measured using Quant-iT PicoGreen assay (Invitrogen) using manufacturer’s protocol.

### 2.13 RNA quantification

MADM OPCs were encapsulated in ELP hydrogels as previously described and cultured for up to 5 days. Hydrogels were collected in 700 μl TRIzol Reagent (Invitrogen), homogenized using a pestle, and sonicated for 10 seconds. RNA was isolated using manufacture’s protocol without modification. 2 μl GlycoBlue was used to facilitate RNA precipitation. RNA purity was confirmed with 260/280 nm absorbance ratio > 1.7 and 260/230 nm absorbance ratio > 1 (BMG LABTECH CLARIOstar monochromatic microplate reader), and the concentration was normalized to 30 μg/ml. Complimentary DNA (cDNA) was made using an iScript cDNA synthesis kit (Bio-Rad, Bio-Rad CFX connect real-time polymerase chain (RT-PCR) machine). PCR was performed using the SsoAdvanced universal SYBR green supermix kit (Bio-Rad, Bio-Rad CFX connect RT-PCR machine). Custom primers were used for gene amplification (Table 2, Bio-Rad)^37,38^.

**Table 2:**
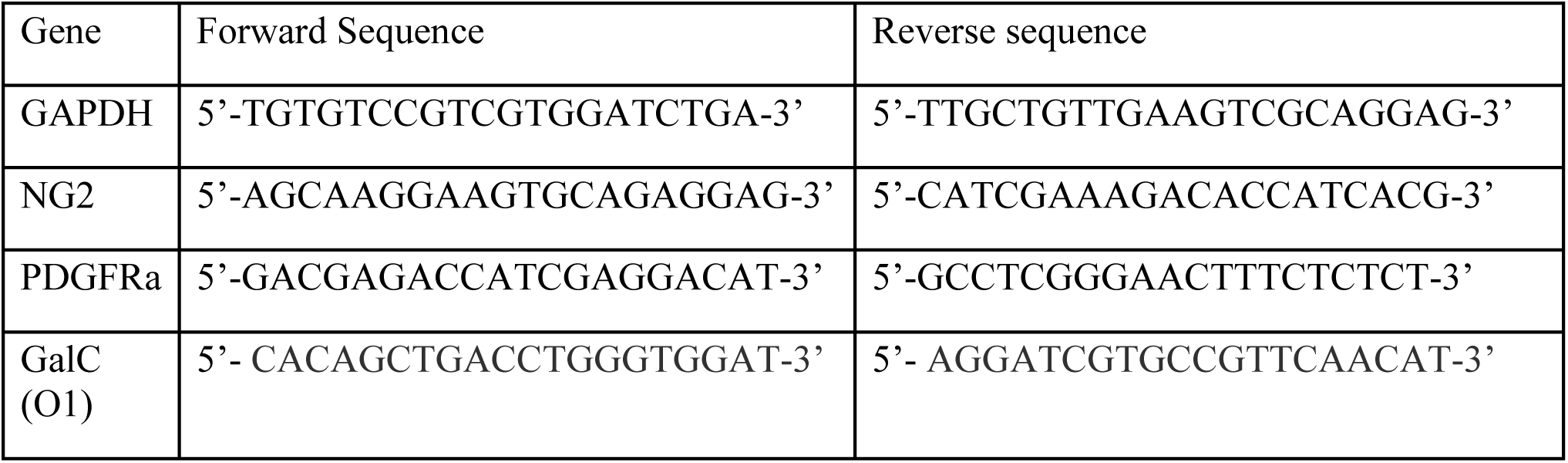
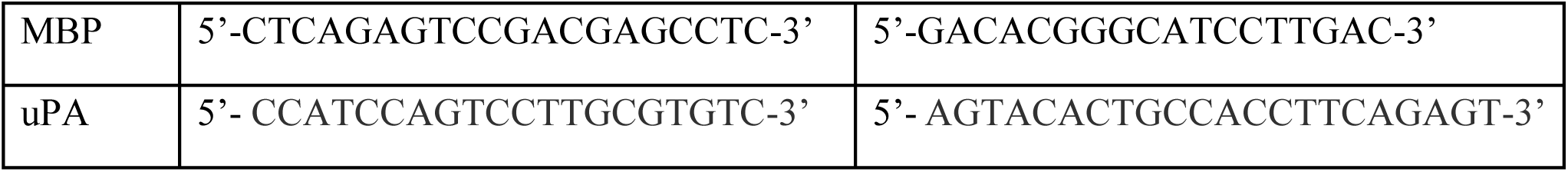
List of primers used for quantitative reverse transcription polymerase chain reaction (RT-qPCR).

### 2.14 Statistical analysis

Data is presented as average ± standard error (SE). Statistical analysis was performed with the student T-test, and significance was determined by p < 0.05.

## 3 Results and Discussion

### 3.1 Azide functionalization of ELPs

Primary amines on each ELP sequence were modified with azides using the diazo transfer reaction (Figure 1)^33,34^. Azide modification was confirmed by the appearance of a peak at 2100 cm^−1^ on the FTIR spectrum, and the intensity of the peak increased with the addition of more azide groups (Supplemental Figure 1). The number of azides attached to each ELP molecule were quantified with high resolution ESI-MS (Supplemental Figure 2). The conversion of each primary amine into an azide group resulted in a 26 Da increase in the protein molecular weight (Mw, Supplemental Figure 2A). One peak appeared in the mass spectra of the pre-modified ELP sequences, confirming that the expression and purification process produced a monodisperse protein (Supplemental Figure 2B-D)^27^. Post diazo transfer reaction, the ELP mass spectra had multiple peaks, confirming that the reaction resulted in a distribution of azide addition (Supplemental Figure 2B-D)^34^. The mass increase of each of these peaks, relative to the unmodified ELP peak, was used to determine the number of azides attached, and the relative peak intensities were used to quantify the azide functionalization distribution. On average 6.28, 3.78, and 3.74 azide groups were added to ELP-u1, ELP-RGD, and ELP-RDG sequences used for MADM OPC encapsulations, respectively. We extended the findings of previous literature here by demonstrating that the average number of azide groups added to a protein can be controlled by stoichiometrically limiting the amount of azide available during the diazo transfer reaction^33,34^.

The addition of azide groups to ELPs decreased their T_t_ (Figure 2A). This occurred because the conversion of amines into azide functional groups increases the hydrophobicity of ELPs^39^. Similar work demonstrated that the opposite effect occurs when ELP methionine residue side chains were modified to become more hydrophilic^40^. In addition, analogous work demonstrated that the ELP T_t_ decreases when the guest residue in the penta-peptide repeating sequence is swapped for a more hydrophobic amino acid^41^. If we consider the azide modification of lysine residues on these ELPs similar to replacing the lysine guest residue with a more hydrophobic amino acid, then a reduction in the T_t_ is the expected result. The T_t_ shift was dependent on the degree of azide functionalization, with the addition of more azide groups resulting in a lower T_t_ (Figure 2B). This is consistent with previous findings that the T_t_ of ELP sequences decreased as the guest amino acid residues are gradually swapped from alanines to more hydrophobic valines^41^ Azide functionalization of ELP-u1, ELP-RGD, and ELP RDG used for MADM OPC encapsulations shifted the T_t_ from 35.6, 35, and 34.3°C to 26.6, 25.5 and 24.3°C, respectively (Figure 2A).

### 3.2 Independent tuning of hydrogel bioactivity and stiffness

Hydrogels were formed by mixing azide-modified ELPs with a PEG-BCN cross-linker using strain-promoted azide alkyne cycloaddition (SPAAC) chemistry (Figure 1). uPA degradable hydrogels were formed by mixing ELP-RGD and ELP-u1 in a 1:1 ratio, and non-degradable hydrogels were formed by mixing ELP-RGD and ELP-RDG in a 1:1 ratio (Figure 1). Rheological time sweeps confirmed previous studies that indicate SPAAC cross-linking occurs on the order of several minutes (Figure 3A)^33,42^. The crossover where the storage modulus (G’) exceeded the loss modulus (G”) indicated that the solution-to-gel transition occurred between 5-6 min for both degradable and non-degradable hydrogels (Figure 3A). The cross-linking reaction was completed within 20 min as indicated by the plateauing of G’ (Figure 3A). These ELP hydrogels have suitable cross-linking rates for cell encapsulation.

**Figure 3:**
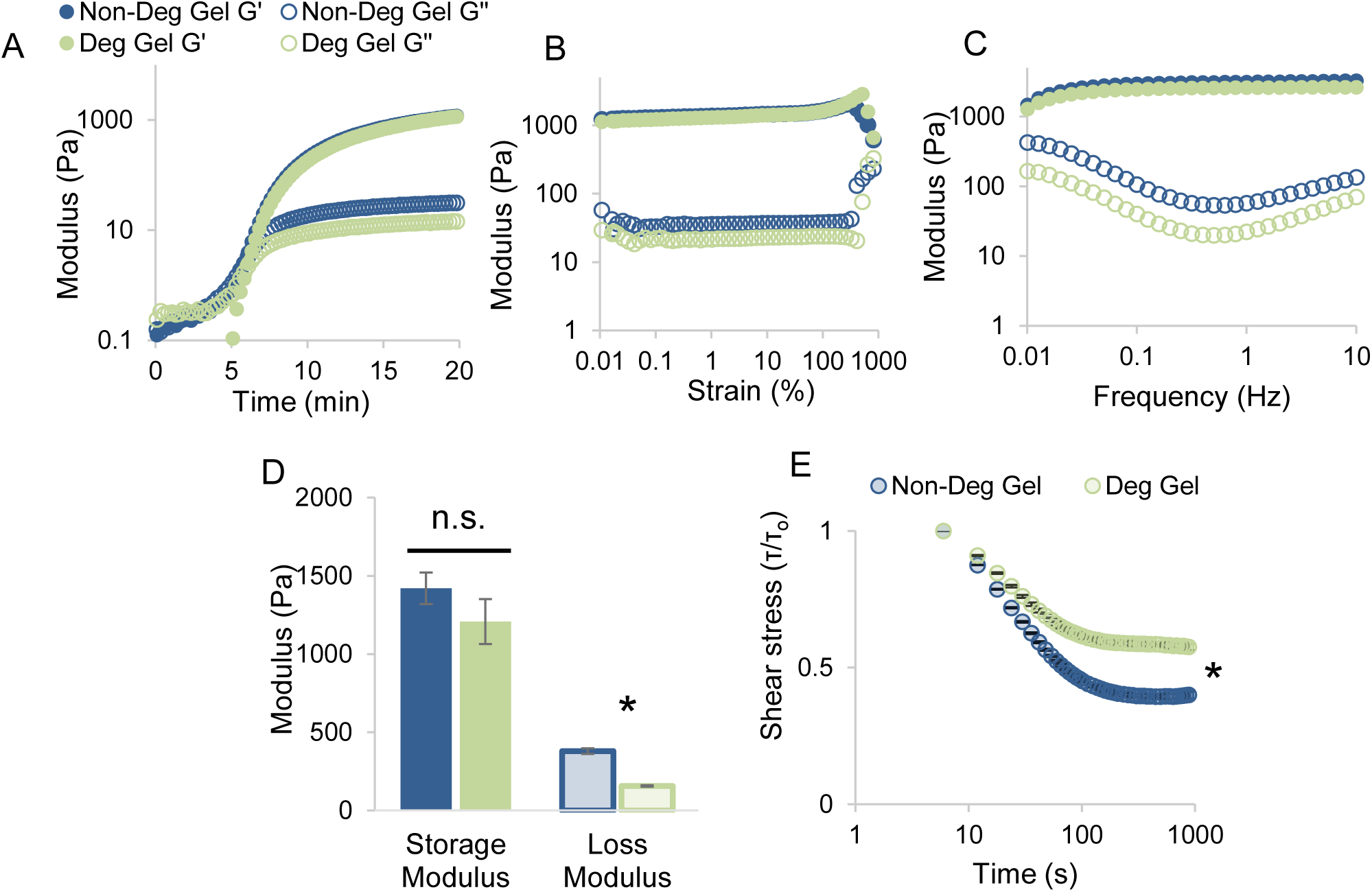
Biomechanics of uPA degradable (Deg) and non-degradable (Non-Deg) ELP hydrogels. (A) Representative time sweeps indicate gelation occurs within 6 min and is complete after 20 min. (B) Representative strain sweeps show the linear viscoelastic range exists up to 300 % strain. (C) Representative frequency sweeps demonstrate hydrogel elastic and loss modulus are frequency dependent. (D) The average storage modulus was similar for both hydrogels while the average loss modulus differed. The storage modulus of both hydrogels was consistent with native neural tissue white matter tracts, where oligodendrocyte precursor cells (OPCs) and oligodendrocytes (OLs) reside. (E) uPA degradable and non-degradable hydrogels relax 43 and 60 % of initial stress over 3.5 min, respectively. Error bars represent standard error (SE) (n ≥ 4). Asterisks indicate statistical significance using student T-test (p < 0.05) and n.s. indicates no statistical significance (p > 0.05).

The similarity between the ELP sequences should create uPA degradable and non-degradable hydrogels with similar biomechanics^26,43^. The linear viscoelastic range (LVER) of both hydrogels was 0 -300 % strain, after which the hydrogel network was irreversibly altered (Figure 3B). Most protein-based hydrogels have a much lower LVER, but dosing PEG into hydrogel network results in a LVER resembling that of PEG-based hydrogels^32,44^. The PEG-BCN crosslinker is large (10 kDa) and the LVER indicated that it impacted the hydrogel biomechanics. The G’ and G” were frequency dependent, which is consistent with previous work (Figure 3C)^33^. The G’ of uPA degradable and non-degradable hydrogels were 1.2 ± 0.14 and 1.42 ± 0.10 kPa (Figure 3D), respectively. These G’ values were not statistically different, confirming that similar hydrogel stiffness was maintained despite the differences in ELP sequences used (Figure 3D). There were differences in the extent of azide functionalization between ELP-u1 and ELP-RDG, which can impact cross-linkability. However, it is likely that for the hydrogels used in this study the differences in ELP azide functionalization did not greatly impact hydrogel biomechanical properties (gelation time and storage modulus) because the cross-linker (PEG-BCN) was the limiting factor in determining the total number of cross-links formed. In addition, by controlling the total ELP concentration and the cross-linking ratio, the G’ of both hydrogels was tuned to be similar to that of native brain tissue white matter tracts (1.895 ± 0.592 kPa), where most native OPCs and OLs reside^45,46^. These ELP hydrogels have biomechanical properties suitable to be *in vitro* mimetics of native brain tissue.

The G” of non-degradable hydrogels was higher than that of uPA degradable hydrogels (Figure 3D). This indicated that non-degradable hydrogels experienced higher viscous losses when strained and translated to a higher degree of stress relaxation (Figure 3E). uPA degradable and non-degradable hydrogels dissipated 43 ± 0.013 % and 60 ± 0.013 % of the initial stress within 3.5 min, respectively, before a plateau was reached (Figure 3E). Stress relaxation rate differences are known to impact encapsulated mesenchymal stem cell spreading, proliferation and differentiation fate^47,48^. The stress relaxation rates of uPA degradable and non-degradable hydrogels, as indicated by the slopes of the curves, were similar to each other (Figure 3E). In addition, the stress relaxation curves for both hydrogels emulate that of brain, liver and adipose tissue^47^. It is unclear how the difference in the magnitude of stress relaxation impacts encapsulated cell behavior. Taken together our results indicated that uPA degradable and non-degradable hydrogels had similar, but not identical, biomechanics that mimic native brain tissue.

### 3.3 MADM OPC uPA expression

We were unsure as to whether or not MADM OPCs expressed uPA because OPC uPA expression *in vivo* is transient, and the p53 and Nf1 gene knockouts may have altered the uPA expression profile of the MADM OPC line^12,30,31^. uPA enzymatic activity of MADM OPCs was determined using a modified zymography technique. Traditionally zymography involves co-cross-linking an enzymatically degradable gel (called the substrate), such as gelatin, alongside a poly(acrylamide) gel^17,21,49,50^. Instead of co-cross-linking ELP alongside poly(acrylamide) we immobilized ELP onto the primary amines readily available on the poly(acrylamide) gel network using THPC, an amine reactive molecule (Supplemental Figure 3A, B). ELP dosed into the poly (acrylamide) gel traveled downwards during electrophoresis, but immobilized ELP remained evenly distributed (Supplemental Figure 3C, D). Given the prevalence of amines in proteins and small peptides, use of this modified zymography technique to determine proteolytic enzymatic activity has extensive utility in the biomaterials field because it expands the number of digestible substrates that can be used. This, in turn, also expands the zymography technique to include exploring the activity of proteolytic enzymes that do not cleave traditionally used substrates like gelatin.

Wells loaded with recombinant uPA and run on a poly(acrylamide) gel dosed with immobilized ELP-u1 confirmed that ELP-u1 was cleavable by both high and low Mw uPA (Figure 4A)^26^. MADM OPCs from a monolayer culture expressed only low Mw uPA, as indicated by a single band that appears adjacent to the low Mw positive control band (Figure 4A). The uPA enzyme exists in two Mw’s, 51.6 and 34.5 kDa, that have similar peptide cleavage enzymatic activity, but high Mw uPA has a higher affinity for activating the plasminogen system^51^. No bands appeared from homogenized MADM OPCs encapsulated in ELP hydrogels (Figure 4A). This could mean that MADM OPCs lose their expression of uPA after being encapsulated in ELP hydrogels. However, it is more likely that this zymography technique was not sensitive enough to detect the low quantities of uPA present; the number of cells in the MADM OPC monolayer were ∼100x that of MADM OPCs encapsulated within the ELP hydrogels, and we were unable to homogenize and concentrate 100 gels into a small enough volume. No band appeared from concentrated cell media, indicating that low Mw uPA was expressed by MADM OPCs and not coming from an external source (Figure 4A).

**Figure 4:**
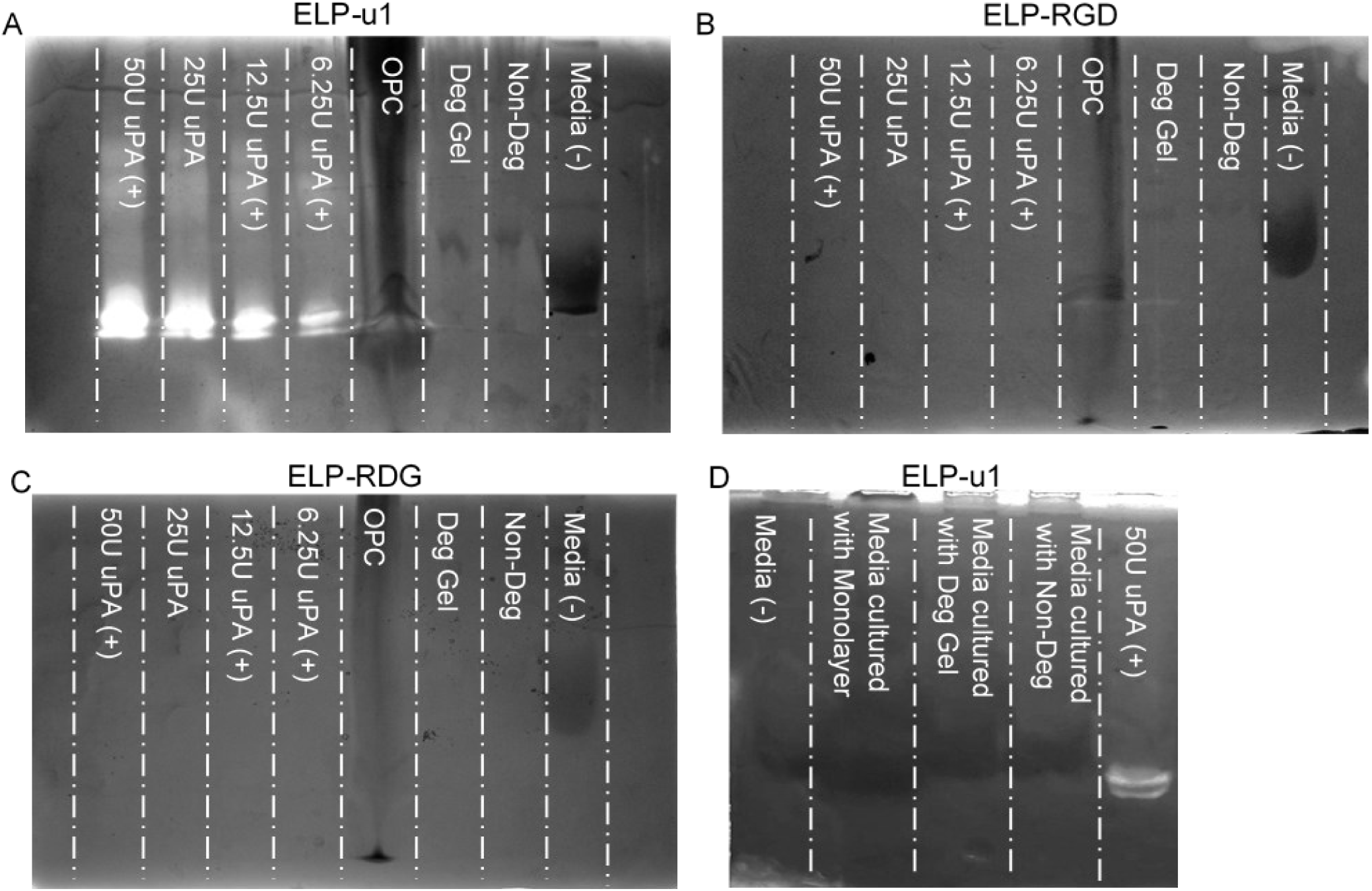
Zymograms of poly(acrylamide) gels dosed with immobilized (A, D) ELP-u1, (B) –RGD, (C) –RDG. (A, B, C) MADM OPCs express low molecular weight uPA to cleave ELP-u1, and do not express any enzymes capable of cleaving ELP-RGD and ELP-RDG sequences. Wells loaded with recombinant uPA (positive control), homogenized MADM OPCs grown in monolayer, homogenized MADM OPCs encapsulated in degradable gel, homogenized MADM OPCs encapsulated in non-degradable gel, and base OPC culture media (negative control). (D) Wells of a gel dosed with immobilized ELP-u1 protein were loaded with OPC media (negative control), recombinant uPa (positive control), and conditioned media from OPCs in monolayers, conditioned media from OPCs encapsulated in uPa-degradable hydrogels, and conditioned media from OPCs encapsulated in non-degradable hydrogels. MADM OPCs do not release soluble uPA into the surrounding media environment; it remains cell membrane bound and only found in zymogram wells

Zymograms of poly(acrylamide) gels dosed with immobilized ELP-RGD and ELP-RDG confirmed that these sequences are not cleavable by uPA (Figure 4B, C)^26^. The MADM OPC monolayer also did not produce any band on these zymograms (Figure 4B, C). This demonstrated that MADM OPCs did not produce any enzymes capable of cleaving ELP-RGD and ELP-RDG. Although these two sequences were not designed to be enzymatically degradable, cells have the machinery to modify peptides and recent work has shown that ELP-RGD is enzymatically cleavable by a disintegrin and metalloprotease 9 (ADAM9)^52^. No bands appeared on zymograms loaded with conditioned media from MADM OPCs (Figure 4D). This confirms that uPA remains bound to the cell membrane in complex with its receptor (uPAR) and is not released into the surrounding environment^11^.

Fluorescent images of the GFP signal expressed by MADM OPCs demonstrated that they degrade uPA degradable hydrogels (Figure 5). Over 3 days, regions with large clusters of cells appeared within the uPA degradable hydrogels. Both the size of the cell clusters and the area of the degraded regions increased over time. We hypothesize that these large cell clusters were a result of cells collapsing onto other cells (along the z-direction) to form multi-layers as the hydrogel matrix was cleaved. Degradation in these hydrogels was heterogeneous and appeared to have a nucleation point. The observed heterogeneous nature of degradation within these hydrogels may be occurring because of the ELP LCST transition. Encapsulated MADM OPCs are incubated at temperatures above the ELP LCST, and the ELP conformational change that occurs while the protein is immobilized within the hydrogel network may cause the uPA-cleavable peptide sequence portion of the protein to be more accessible to MADM OPCs in some regions than in other regions. In non-degradable hydrogels, MADM OPCs remained homogenously distributed in small, isolated clusters over the 3 days (Figure 5). This type of growth was similar to growth patterns previously observed when MADM OPCs were encapsulated in non-degradable PEG hydrogels^53^. Taken together, this data demonstrated that MADM OPCs were able to degrade uPA degradable hydrogels because of their expression of low Mw uPA, and they were unable to remodel the non-degradable ELP hydrogel matrix.

**Figure 5:**
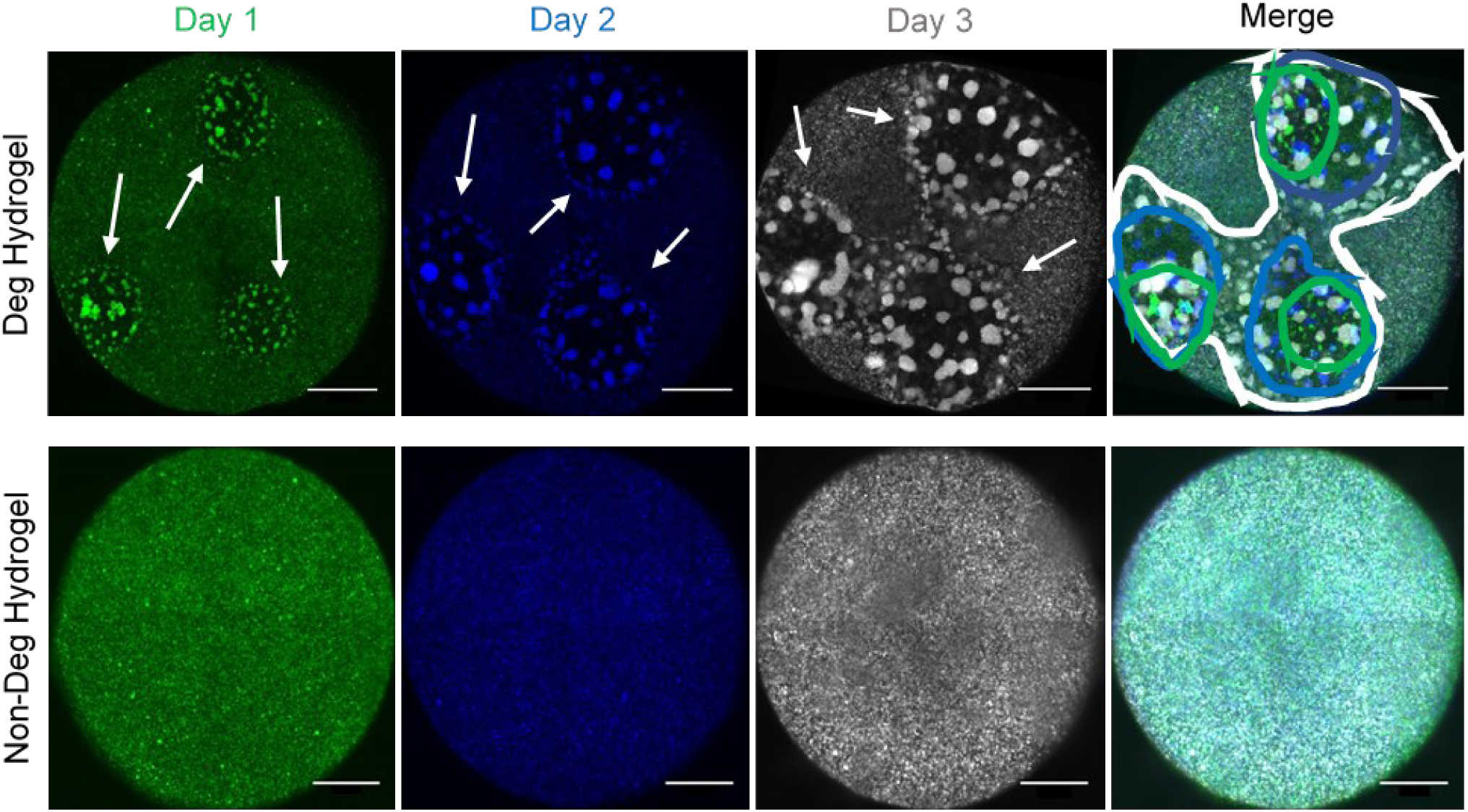
MADM OPC GFP+ signal (represented in green for day 1, blue for day 2, and gray for day 3) of representative ELP hydrogels with and without uPA cleavage sites over 3 days. Hydrogels with uPA cleavage sites have regions with visible degradation that increase in area over 3 days. Degradation is not observed in non-degradable hydrogels. Lines in merged degradable hydrogel image display area of degraded region over time; green, blue and white represent day 1, 2, and 3, respectively. Scale bars are 1,000 μm.

### 3.4 Encapsulated MADM OPC metabolic activity

Encapsulated MADM OPCs had similar viability in degradable and non-degradable hydrogels, with 77 ± 3.6 and 80 ± 2.3 % of cells remaining viable after 3 days, respectively (Figure 6A, B). The viability of MADM OPCs was on the lower end of what is typically observed for cells encapsulated within ELP hydrogels^27,33,54^. We attribute the lower viability to the high sensitivity of MADM OPCs towards cross-linking chemistry. MADM OPCs encapsulated in ELPs cross-linked with amine reactive THPC, which is cytocompatible with several cell types^27,54^, were not viable after gelation (data not shown). SPAAC chemistry is considered bio-orthogonal, meaning it does not cross-react with cellular machinery, however, there could be some reactivity of the strained alkyne bond with cellular machinery^55^. The increase in ATP and DNA content of MADM OPCs encapsulated over time confirmed that MADM OPCs are viable and proliferative in ELP hydrogels cross-linked using SPAAC chemistry (Figure 6C, D).

**Figure 6:**
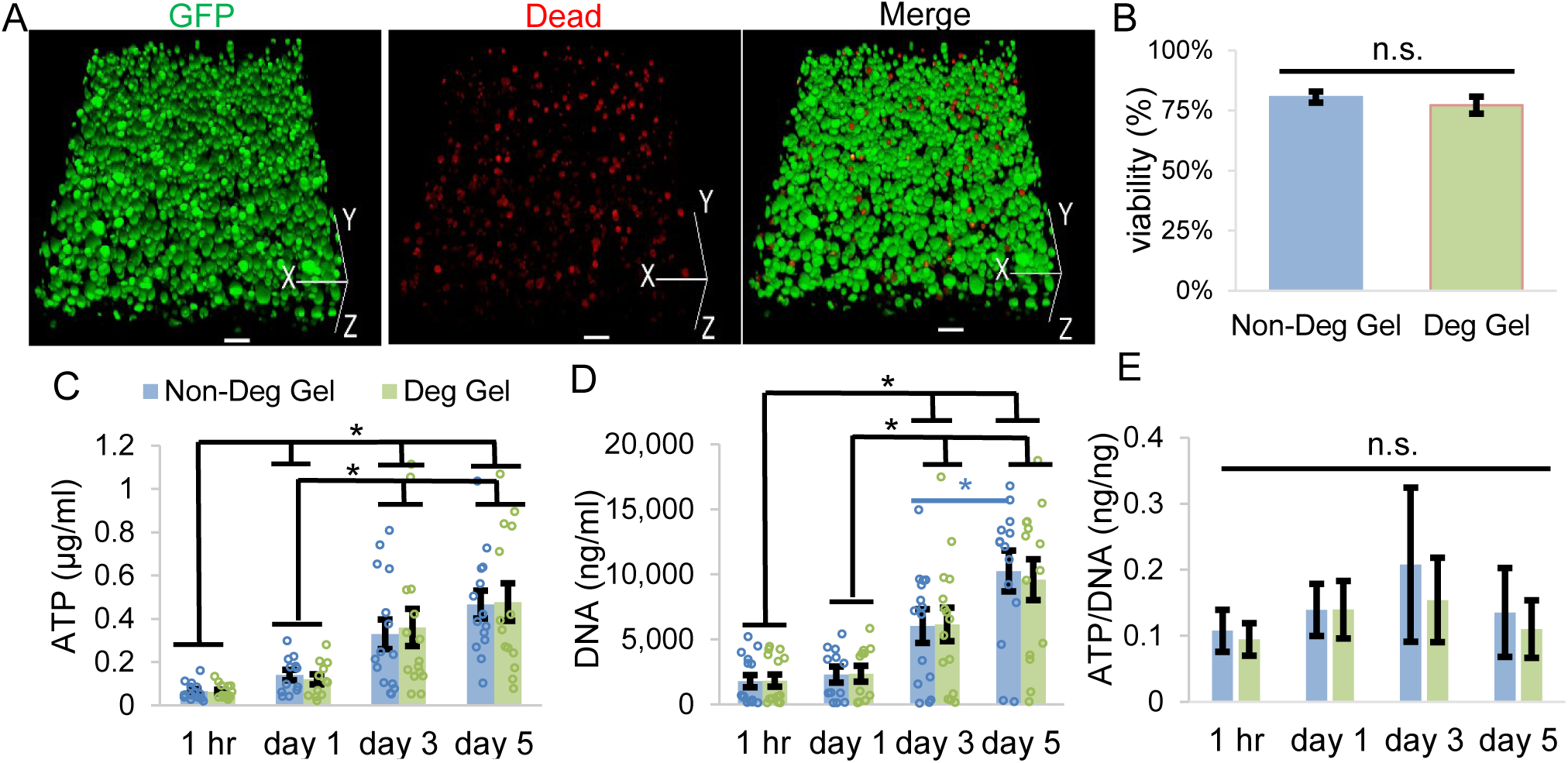
Encapsulated MADM OPC viability and metabolic activity. (A) Representative 3D-projection (252 μm z-stack) of MADM OPCs encapsulated and cultured in uPA degradable hydrogel for 3 days (green is GFP, red is dead cells). Scale bars are 100 μm. (B) The viability of MADM OPCs encapsulated in ELP hydrogels was not affected by the uPA-degradability of the hydrogel matrix. Error bars represent SE (n = 3). Encapsulated MADM OPC (C) ATP concentration, (D) DNA concentration, and (E) ATP/DNA ratio was similar in both uPA degradable and non-degradable ELP hydrogels over 5 days. Error bars represent SE (n ≥ 12). Asterisks indicate statistical significance using student T-test (p < 0.05) and n.s. indicates no statistical significance (p > 0.05).

MADM OPC metabolic activity was similar in both hydrogel systems over 5 days (Figure 6C, D, E). While the total ATP and DNA content per hydrogel increased, on a per cells basis (ATP/DNA) there was no statistically significant shift in the metabolic activity over time (Figure 6C, D, E). In addition, no statistically significant differences in metabolic activity developed between MADM OPCs encapsulated in uPA degradable hydrogels versus those encapsulated in non-degradable hydrogels (Figure 6C, E). The differentiation of OPCs to OLs is marked by a decrease in cellular metabolic activity because OLs are not proliferative while OPCs are^56^. ATP and DNA data did not indicate that uPA degradable hydrogels impacted MADM OPC maturation. MADM OPC proliferation was also similar in both uPA degradable and non-degradable hydrogels (Figure 7). While slightly less MADM OPCs were proliferative in uPA degradable hydrogels (33.2 ± 2.8 %) versus non-degradable hydrogels (36.6 ± 3.5 %) on day 3, the difference was not statistically significant (Figure 7B). Taken together, the data suggests that MADM OPC viability and metabolic activity was not influenced by the presence of uPA degradability in ELP hydrogels.

**Figure 7:**
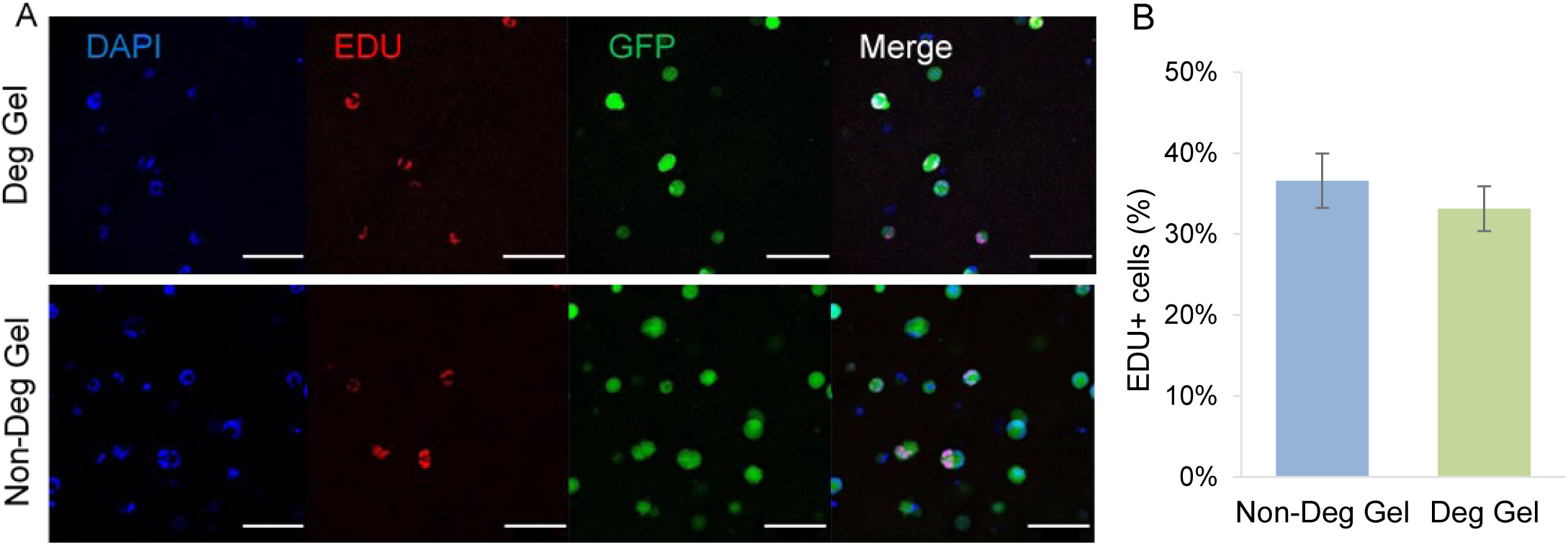
Encapsulated MADM OPC proliferation at day 3 of culture. (A) Representative z-projections of 20 μm stacks (blue is DAPI, red is EdU, green is GFP). Scale bars are 100 μm. (B) A similar fraction of encapsulated MADM OPCs were proliferative in uPA degradable and non-degradable hydrogels after 3 days. Error bars represent SE (n = 3). n.s. indicates no statistical significance using student T-test (p > 0.05).

### 3.5 MADM OPC maturation in ELP hydrogels

OPC maturation to OLs is measured through a morphological increase in process extensions and branching^14^. Encapsulated MADM OPC morphology was assessed by labeling f-actin (Figure 8). Some encapsulated MADM OPCs extended processes within both uPA degradable and non-degradable hydrogels, but this was limited to cells near the surface of the hydrogels (Figure 8A). The vast majority of MADM OPCs, especially ones located deeper into the hydrogel matrix, grew in circular clusters (Figure 8A), similar to previous work^53^. Encapsulated MADM OPC morphology was not indicative of maturation in either hydrogel.

**Figure 8:**
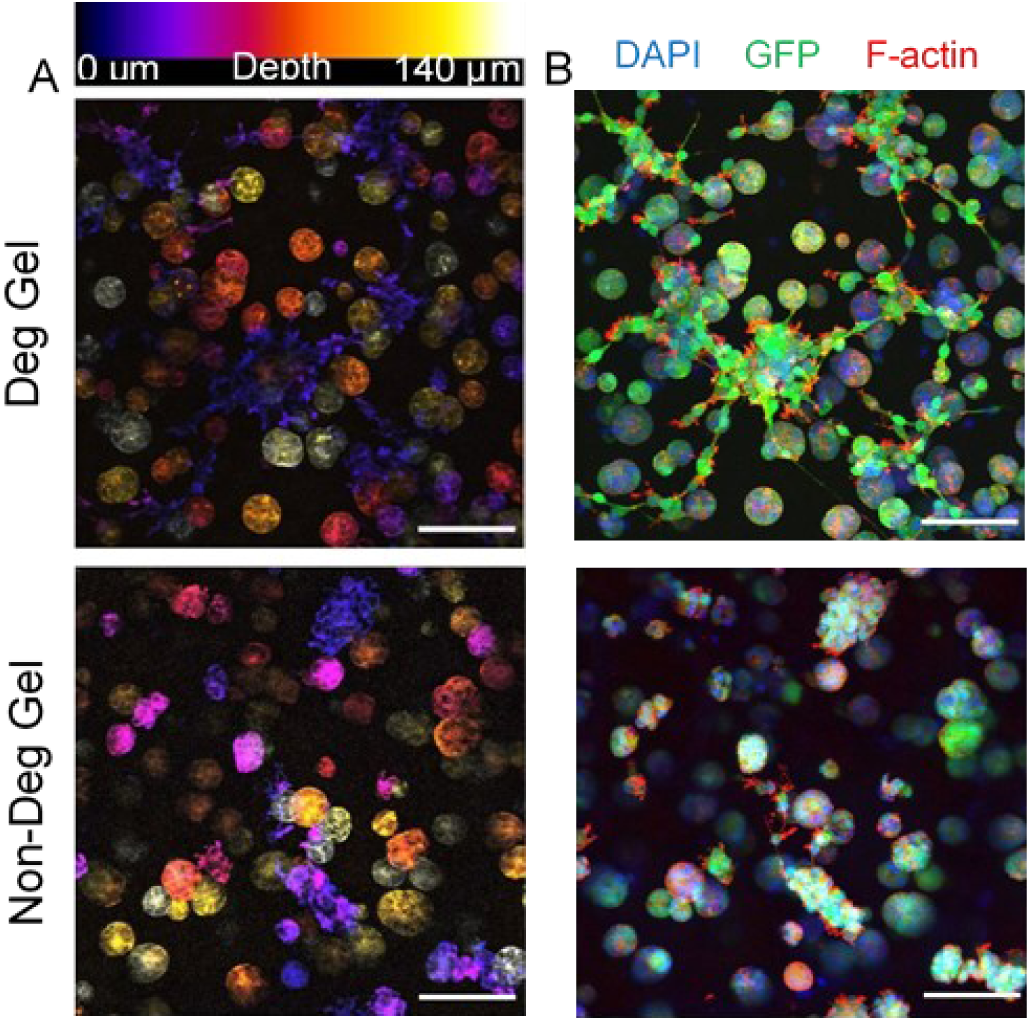
Encapsulated MADM OPC morphology. (A) Confocal depth projections display MADM OPC F-actin expression within uPA degradable and non-degradable hydrogels cultured for 4 days. The color indicates the depth from the hydrogel surface (with orange to yellow being deepest into the hydrogel). Encapsulated MADM OPCs appear to only develop significant processes near the surface of ELP hydrogels. (B) Max projection of the stacks shown in Figure 8A with blue staining for DAPI, green representing GFP, and red staining for phalloidin. Scale bars are 100 μm.

Several gene and protein expression changes occur during OPC maturation^57–61^. We examined encapsulated MADM OPC expression of four genes relevant to the maturation process (Figure 9A)^60,61^. Neural antigen 2 (NG2) and platelet derived growth factor receptor alpha (PDGFRα) are genes expressed by OPCs and their expression decreases as they mature into OLs. The gene expression of these early OPC markers decreased over 5 days in MADM OPCs encapsulated in degradable hydrogels relative to non-degradable hydrogels (Figure 9B). NG2 gene expression was higher in uPA degradable hydrogels at day 1, but then decreased from day 1 to day 3. PDGFRα was lower on day 1 in uPA degradable hydrogels and continued to decrease over 5 days. The gene expression of PDGFRα was lower than that of NG2 on day 5, which agrees with previous work that NG2 is expressed up until the immature OL stage, while PDGFRα is exclusively expressed only by OPCs^60,61^. This data indicated that encapsulated MADM OPCs were moving towards a more mature OPC state in uPA degradable hydrogels.

**Figure 9:**
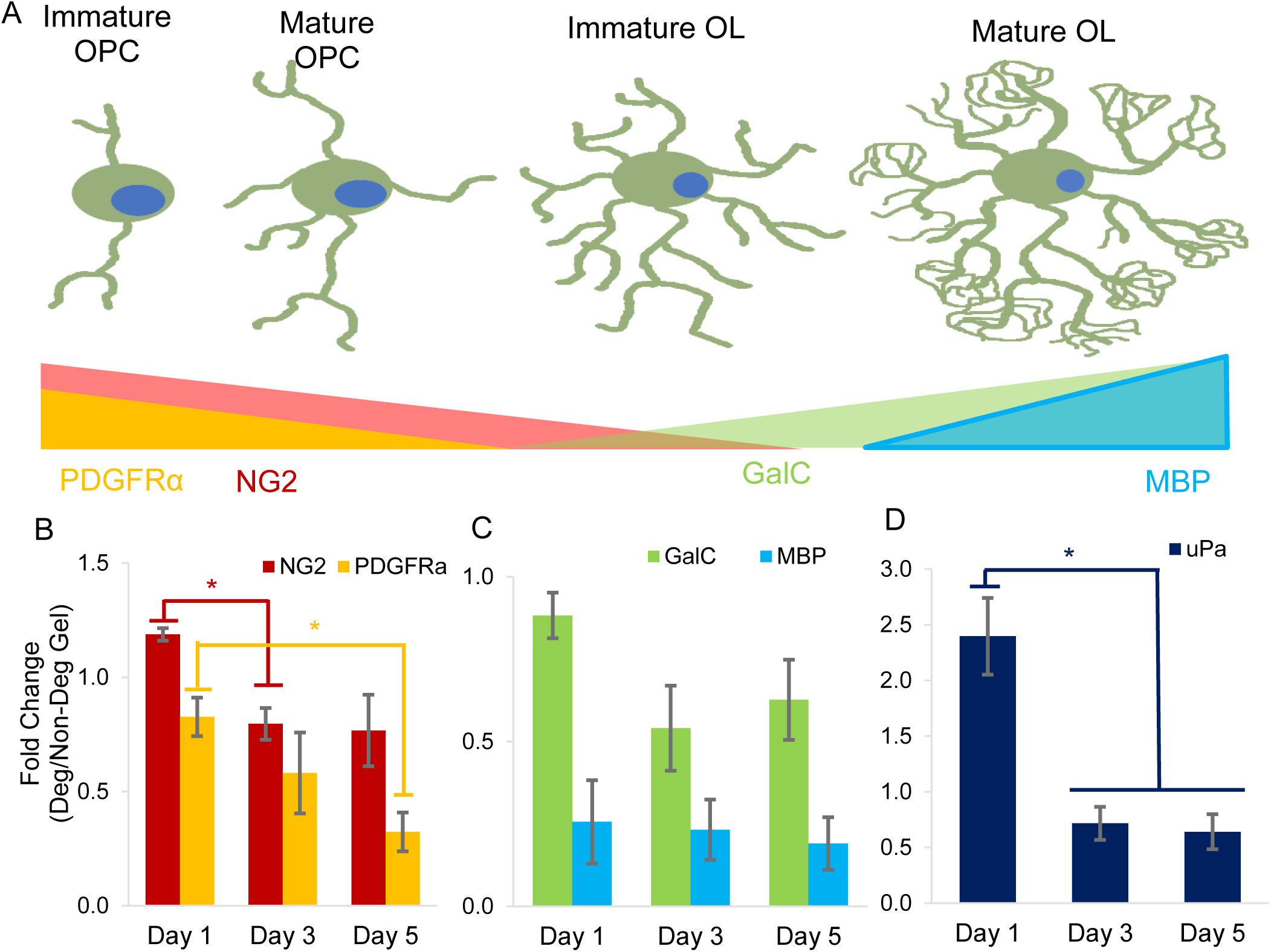
Encapsulated MADM OPC gene expression over 5 days. (A) As OPCs differentiate into oligodendrocytes and oligodendrocytes mature, NG2 and PDGFRα gene expression is reduced, while GalC and MBP gene expression increases. (B) Within our 3D hydrogel system, MADM OPC gene expression of NG2 and PDGFRα decreased over time in uPA degradable hydrogels relative to non-degradable hydrogels, indicating that uPA degradation promotes maturation of immature OPCs. (C) MADM OPC gene expression of GalC and MBP do not increase over time when encapsulated in uPA degradable hydrogels relative to non-degradable hydrogels. This indicated that uPA degradation does not induce the fate transition from OPC to OL. (D) uPA gene expression is initially upregulated for MADM OPCs encapsulated in uPA degradable hydrogels before dropping to levels below that of MADM OPCs encapsulated in non-degradable hydrogels. Error bars represent SE (n = 3). Asterisks indicate statistical significance using student T-test (p < 0.05).

Expression of the galactosylceramidase (GalC) protein is an identifier for when an OPC has transitioned into an OL, and GalC gene expression precedes this in a similar manner (Figure 9A). Increased expression of myelin basic protein (MBP) gene identifies the transition from an immature OL to a myelinating one (Figure 9A). Gene expression of GalC and MBP was higher in MADM OPCs encapsulated in non-degradable hydrogels than those encapsulated in uPA degradable hydrogels (Figure 9C). The gene expression of GalC decreased in uPA degradable hydrogels relative to non-degradable hydrogels from day 1 to day 3. MBP gene expression remained static over the 5 days. If MADM OPCs were maturing into OLs, the gene expression of these two markers should have increased over time. Instead MBP is constant and GalC decreases over time. In addition, this data disagrees with previous research that indicates OPCs do not express GalC and MBP^60^. Taken together this data indicates that uPA degradable hydrogels promote MADM OPC maturation, but do not induce the transition from OPC to OL.

Based on all data (morphology, metabolic activity, and OPC gene expression) MADM OPCs encapsulated in uPA degradable hydrogels were not transitioning from OPCs to OLs. The lack of MADM OPC maturation into myelinating OLs could be due to the cell line itself. MADM OPC differentiation *in vivo* is delayed relative to normal development^31^. This is likely caused by disruptions to the Erk 1/2 and mTORC1 pathways that result from the Nf1 and p53 gene knockouts (Supplemental Figure 4). In particular, p53 inhibits Akt, which digests Mek 1/2, and activates PTEN, which inhibits PI3K (Supplemental Figure 4). The knockout of the p53 gene does not allow for Erk 1/2 to activate and prevents the inhibition of the mTORC1 pathway (Supplemental Figure 4). Pharmacological disruptions to the Erk 1/2 and mTORC1 pathways inhibited OPC maturation to OLs *in vitro* and delayed it *in vivo*^59,62,63^. It is not known what trophic support OPCs receive *in vivo* to overcome disruptions to the Erk 1/2 and mTORC1 pathways. Previous literature indicates that MADM OPC maturation within ELP hydrogels could be affected by disruptions to the Erk 1/2 and mTORC1 pathways.

The zymography technique did not have the sensitivity necessary to quantify uPA enzymatic activity changes from the encapsulation process with densitometry (Figure 4). To determine if encapsulating MADM OPCs in uPA degradable hydrogels affected their expression of uPA, uPA RNA content was measured over 5 days (Figure 9D). Encapsulated MADM OPC uPA gene expression was higher in uPA degradable hydrogels relative to non-degradable hydrogels on day 1 (Figure 9D). Then the uPA gene expression of MADM OPCs encapsulated in degradable hydrogels dropped below that of MADM OPCs in non-degradable hydrogels on days 3 and 5 (Figure 9D). The rapid initial increase in uPA gene expression that was followed by inhibition could be a result of MADM OPCs becoming saturated with uPA after an initial upregulation of transcription. These data demonstrated that MADM OPCs do upregulate uPA gene expression when exposed to uPA degradable hydrogels.

This work demonstrates that the incorporation of enzymatic degradability into hydrogel networks can be used to do more than degrade the material, it can be used to influence cell behavior and fate. Previous literature found that neural progenitor cell (NPC) stemness was maintained within enzymatically degradable hydrogels, and that degradation primed cells for differentiation^52^. uPA degradable hydrogels may have impacted MADM OPCs in a similar manner. MADM OPCs were primed for differentiation by transitioning to a more mature OPC, but maintained their stemness because they never transitioned from OPCs to OLs (Figure 9). However, we argue that the specific enzyme(s) used to degrade the hydrogel matrix matters, with different enzymes leading to different cellular outcomes. uPA degradable hydrogels promoted MADM OPC maturation and disruptions to the Erk 1/2 and mTORC1 pathways prevented further differentiation of this cell line (Figure 9, Supplemental Figure 4). Hydrogel degradation can be utilized to allow encapsulated cells to do more than remodel the matrix; it can be used to curate encapsulated cell behavior.

### 3.6 Impact of encapsulation process on MADM OPC gene expression

We investigated the impact of the encapsulation process on MADM OPC gene expression because the general effects of the encapsulation process on cell behavior are not well studied (Figure 10). Encapsulation of MADM OPCs in both hydrogels greatly increased NG2 gene expression relative to MADM OPCs in monolayer cultures on day 1, and while encapsulated MADM OPC NG2 gene expression decreased over time, it never returned to levels observed in MADM OPCs cultured in a monolayer (Figure 10A). Gene expression of PDGFRα and GalC followed a similar trend, except the initial gene expression increase was mild, and by day 5 expression fell below levels of MADM OPCs cultured in a monolayer (Figure 10B, C). MBP gene expression was found to be different in MADM OPCs encapsulated in degradable and non-degradable hydrogels (Figure 10D). In degradable hydrogels MADM OPC MBP gene expression started out higher than MADM OPCs in monolayer cultures and proceeded to decrease to sub-monolayer culture levels overtime (Figure 10D). In non-degradable hydrogels, MADM OPC MBP gene expression increased substantially relative to MADM OPCs in monolayer culture on day 1, dropped below monolayer levels on day 3, and then increased again on day 5 (Figure 10D). MADM OPC uPA expression was also different in the two hydrogel systems. In uPA degradable hydrogels MADM OPC uPA gene expression was upregulated relative to monolayer cultures initially and then dropped to sub monolayer levels by day 5 (Figure 10E). In non-degradable hydrogels MADM OPC uPA gene expression was unaffected initially, then increased briefly on day 3 before returning to monolayer levels on day 5 (Figure 10E).

**Figure 10:**
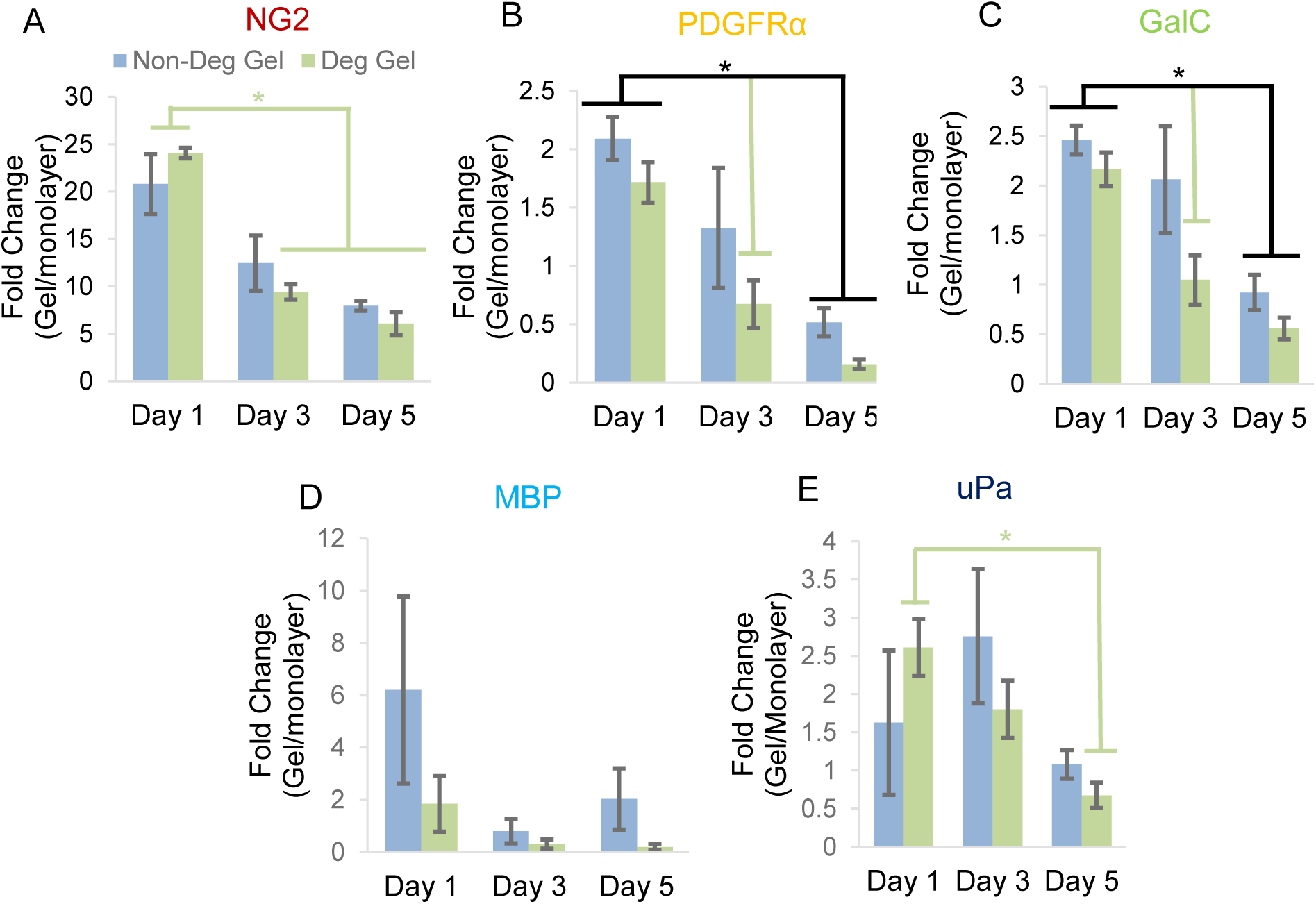
Effect of encapsulation process on MADM OPC gene expression. MADM OPC gene expression of (A) NG2, (B) PDGFRα, (C) GalC, (D) MBP, and (E) uPA was affected by the encapsulation process. Displaying SE (n = 3). Asterisks indicate statistical significance using student T-test (p < 0.05).

The encapsulation process results in large changes to the MADM OPC extracellular environment, and it is difficult to assess why the observed gene expression changes occurred. The results could be from biomechanical changes like stiffness differences between the ELP hydrogels and the tissue culture polystyrene (TCPS) on which monolayers are grown, or the introduction of a 3-dimensional mesh to the MADM OPC environment^9,22^. These gene expression changes could also be induced by bioactivity differences, relative to the plate-based culture system, such as the RGD integrin binding that was incorporated into both uPA degradable and non-degradable ELP hydrogels. We archived the results here to help elucidate the effects of the encapsulation process on gene expression and to inform future studies.

## 4 Conclusions

3D ELP hydrogels can be designed with similar biomechanical properties, to mimic native CNS ECM, and varying bioactivity (i.e., uPA enzymatic degradability) to influence encapsulated cell behavior. Azide functionalization of ELP made the protein more hydrophobic and reduced the T_t_. The average number of azides added to ELP could be controlled by limiting the azide present during the diazotransfer reaction. Three similar ELP sequences were mixed to form hydrogels with independent control over hydrogel stiffness and uPA enzymatic degradation. However, the small differences in the ELP amino acid sequences created hydrogels with different stress relaxation properties. A modified zymography technique was used to prove that MADM OPCs express low Mw uPA and were capable of cleaving the uPA degradable ELP sequence. This modified zymography technique expands the number of digestible substrates that can be used in gel electrophoresis to include any protein or small peptide sequence. MADM OPC viability, metabolic activity and proliferation was not altered by the presence of uPA enzymatic degradation in the ELP hydrogel network. MADM OPCs located near the surface of ELP hydrogels extended processes, but most cells grew in circular clusters deeper in the hydrogels. Gene expression changes in encapsulated MADM OPCs indicated that uPA degradable hydrogels promoted the transition from immature to mature OPCs, but did not promote the transition fully into OLs, likely inhibited by the genetic nature of the MADM OPCs. We demonstrated that enzymatic degradation in hydrogel systems can be used to influence cell behavior post encapsulation in addition to breaking down the hydrogel network.

## 5 Supporting Information

ELP azide functionalization creates the appearance of a peak at 2100 cm^−1^ on FTIR spectra (Supplemental Figure 1). The molecular distribution of azide addition on ELP is quantified with high resolution ESI-MS (Supplemental Figure 2). ELP is immobilized onto the polyacrylamide gel network using amine reactive THPC for zymography experiments (Supplemental Figure 3). Schematic of the Erk 1/2 and mTORC1 pathways and their relationship with the Nf1 and p53 genes (Supplemental Figure 4). This material is available free of charge via the internet at http://pub.acs.org.

## 6 Author Contributions

The manuscript was written through contributions of all authors. All authors have given approval to the final version of the manuscript.

## 8 Acknowledgements

We gratefully acknowledge financial support from NSF grant 1904198 and the Translational Health Research Institute of Virginia for a career development award to KJL. We acknowledge several facilities for providing equipment and expertise: the Keck Center for Cellular Imaging at UVA for the usage of the Zeiss 780 confocal microscopy system (NIH OD016446), the UVA Biomolecular Magnetic Resonance Facility for use of the 500 MHz HNMR system, the UVA Center for Advancing Biomanufacturing for use of the Anton Paar MCR 302 rheometer, the UVA Biomolecular Analysis Facility for collecting MS data, the Giri Lab for use of the FTIR spectrometer, and the Cai lab for use of the Leica SP8 X confocal microscopy system. We also acknowledge Anne Katherine Brooks for her assistance in developing the SPAAC cross-linking hydrogel system.

