## Supplemental information for "Guiding oligodendrocyte precursor cell maturation with urokinase plasminogen activator-degradable elastin-like protein hydrogels"

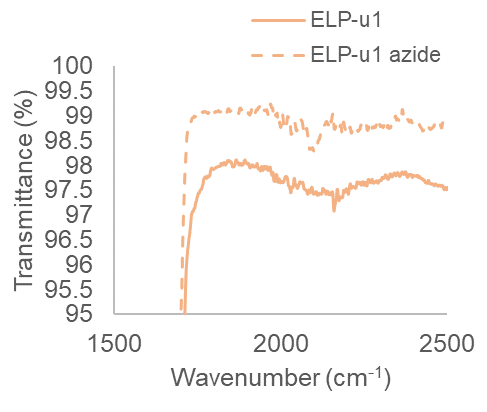

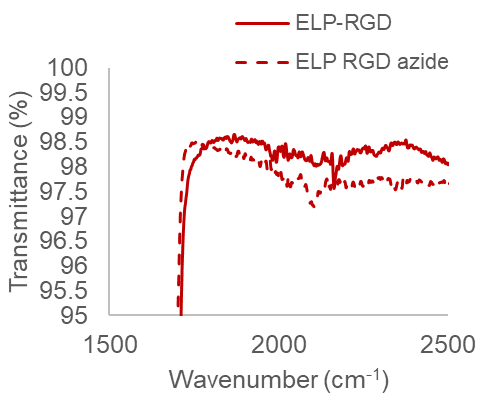

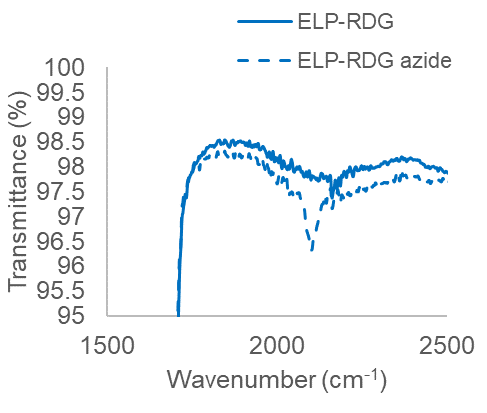

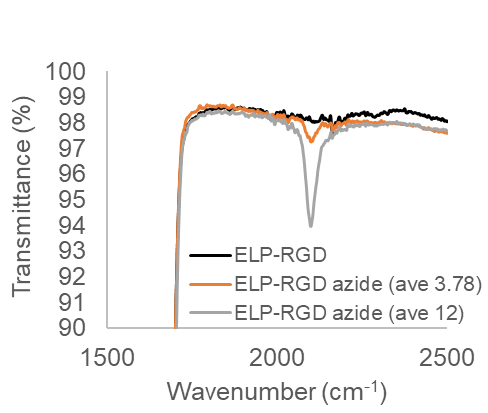

D

A

B

C

Transmittance (%)

| Wavenumber (cm-1) | ELP-u1 | ELP-u1 azide |
| --- | --- | --- |
| 4000 | 98.66 | 99.35 |
| 3999 | 98.66 | 99.36 |
| 3998 | 98.66 | 99.37 |
| 3997 | 98.66 | 99.38 |
| 3996 | 98.66 | 99.39 |
| 3995 | 98.65 | 99.39 |
| 3994 | 98.64 | 99.39 |
| 3993 | 98.64 | 99.4 |
| 3992 | 98.63 | 99.4 |
| 3991 | 98.63 | 99.39 |
| 3990 | 98.63 | 99.39 |
| 3989 | 98.63 | 99.39 |
| 3988 | 98.62 | 99.39 |
| 3987 | 98.62 | 99.39 |
| 3986 | 98.62 | 99.38 |
| 3985 | 98.62 | 99.37 |
| 3984 | 98.64 | 99.35 |
| 3983 | 98.66 | 99.34 |
| 3982 | 98.67 | 99.33 |
| 3981 | 98.68 | 99.34 |
| 3980 | 98.68 | 99.36 |
| 3979 | 98.68 | 99.38 |
| 3978 | 98.67 | 99.4 |
| 3977 | 98.66 | 99.41 |
| 3976 | 98.66 | 99.42 |
| 3975 | 98.66 | 99.42 |
| 3974 | 98.66 | 99.41 |
| 3973 | 98.66 | 99.4 |
| 3972 | 98.65 | 99.39 |
| 3971 | 98.64 | 99.36 |
| 3970 | 98.63 | 99.34 |
| 3969 | 98.63 | 99.32 |
| 3968 | 98.64 | 99.31 |
| 3967 | 98.64 | 99.31 |
| 3966 | 98.64 | 99.32 |
| 3965 | 98.64 | 99.34 |
| 3964 | 98.65 | 99.35 |
| 3963 | 98.65 | 99.37 |
| 3962 | 98.65 | 99.38 |
| 3961 | 98.65 | 99.38 |
| 3960 | 98.65 | 99.38 |
| 3959 | 98.65 | 99.37 |
| 3958 | 98.64 | 99.37 |
| 3957 | 98.64 | 99.37 |
| 3956 | 98.63 | 99.36 |
| 3955 | 98.63 | 99.35 |
| 3954 | 98.64 | 99.34 |
| 3953 | 98.65 | 99.34 |
| 3952 | 98.66 | 99.34 |
| 3951 | 98.66 | 99.33 |
| 3950 | 98.66 | 99.32 |
| 3949 | 98.65 | 99.32 |
| 3948 | 98.65 | 99.32 |
| 3947 | 98.65 | 99.33 |
| 3946 | 98.66 | 99.34 |
| 3945 | 98.67 | 99.34 |
| 3944 | 98.69 | 99.34 |
| 3943 | 98.71 | 99.33 |
| 3942 | 98.72 | 99.32 |
| 3941 | 98.72 | 99.31 |
| 3940 | 98.72 | 99.31 |
| 3939 | 98.72 | 99.32 |
| 3938 | 98.71 | 99.33 |
| 3937 | 98.71 | 99.34 |
| 3936 | 98.71 | 99.35 |
| 3935 | 98.71 | 99.34 |
| 3934 | 98.72 | 99.33 |
| 3933 | 98.72 | 99.31 |
| 3932 | 98.72 | 99.31 |
| 3931 | 98.71 | 99.31 |
| 3930 | 98.7 | 99.32 |
| 3929 | 98.71 | 99.34 |
| 3928 | 98.72 | 99.35 |
| 3927 | 98.73 | 99.35 |
| 3926 | 98.73 | 99.35 |
| 3925 | 98.72 | 99.35 |
| 3924 | 98.71 | 99.35 |
| 3923 | 98.71 | 99.35 |
| 3922 | 98.7 | 99.34 |
| 3921 | 98.7 | 99.34 |
| 3920 | 98.7 | 99.32 |
| 3919 | 98.7 | 99.31 |
| 3918 | 98.71 | 99.3 |
| 3917 | 98.72 | 99.3 |
| 3916 | 98.73 | 99.31 |
| 3915 | 98.73 | 99.32 |
| 3914 | 98.72 | 99.34 |
| 3913 | 98.71 | 99.35 |
| 3912 | 98.71 | 99.36 |
| 3911 | 98.72 | 99.35 |
| 3910 | 98.73 | 99.33 |
| 3909 | 98.75 | 99.32 |
| 3908 | 98.76 | 99.3 |
| 3907 | 98.75 | 99.3 |
| 3906 | 98.75 | 99.3 |
| 3905 | 98.74 | 99.3 |
| 3904 | 98.76 | 99.31 |
| 3903 | 98.77 | 99.31 |
| 3902 | 98.79 | 99.31 |
| 3901 | 98.78 | 99.32 |
| 3900 | 98.77 | 99.34 |
| 3899 | 98.75 | 99.35 |
| 3898 | 98.74 | 99.36 |
| 3897 | 98.73 | 99.36 |
| 3896 | 98.73 | 99.36 |
| 3895 | 98.73 | 99.35 |
| 3894 | 98.73 | 99.35 |
| 3893 | 98.73 | 99.34 |
| 3892 | 98.72 | 99.34 |
| 3891 | 98.72 | 99.33 |
| 3890 | 98.73 | 99.32 |
| 3889 | 98.73 | 99.31 |
| 3888 | 98.73 | 99.3 |
| 3887 | 98.73 | 99.29 |
| 3886 | 98.74 | 99.29 |
| 3885 | 98.75 | 99.28 |
| 3884 | 98.76 | 99.29 |
| 3883 | 98.75 | 99.3 |
| 3882 | 98.74 | 99.32 |
| 3881 | 98.73 | 99.33 |
| 3880 | 98.74 | 99.33 |
| 3879 | 98.75 | 99.32 |
| 3878 | 98.75 | 99.31 |
| 3877 | 98.74 | 99.29 |
| 3876 | 98.73 | 99.28 |
| 3875 | 98.75 | 99.28 |
| 3874 | 98.77 | 99.29 |
| 3873 | 98.8 | 99.29 |
| 3872 | 98.8 | 99.29 |
| 3871 | 98.8 | 99.28 |
| 3870 | 98.8 | 99.28 |
| 3869 | 98.81 | 99.29 |
| 3868 | 98.8 | 99.29 |
| 3867 | 98.78 | 99.28 |
| 3866 | 98.77 | 99.28 |
| 3865 | 98.76 | 99.28 |
| 3864 | 98.76 | 99.28 |
| 3863 | 98.76 | 99.29 |
| 3862 | 98.75 | 99.29 |
| 3861 | 98.75 | 99.3 |
| 3860 | 98.75 | 99.3 |
| 3859 | 98.77 | 99.3 |
| 3858 | 98.81 | 99.29 |
| 3857 | 98.83 | 99.29 |
| 3856 | 98.83 | 99.29 |
| 3855 | 98.8 | 99.3 |
| 3854 | 98.79 | 99.3 |
| 3853 | 98.79 | 99.29 |
| 3852 | 98.8 | 99.28 |
| 3851 | 98.79 | 99.3 |
| 3850 | 98.77 | 99.31 |
| 3849 | 98.76 | 99.34 |
| 3848 | 98.76 | 99.35 |
| 3847 | 98.77 | 99.36 |
| 3846 | 98.77 | 99.36 |
| 3845 | 98.78 | 99.35 |
| 3844 | 98.77 | 99.35 |
| 3843 | 98.76 | 99.36 |
| 3842 | 98.76 | 99.36 |
| 3841 | 98.77 | 99.35 |
| 3840 | 98.77 | 99.34 |
| 3839 | 98.77 | 99.33 |
| 3838 | 98.77 | 99.32 |
| 3837 | 98.77 | 99.32 |
| 3836 | 98.77 | 99.33 |
| 3835 | 98.77 | 99.33 |
| 3834 | 98.77 | 99.33 |
| 3833 | 98.76 | 99.34 |
| 3832 | 98.75 | 99.35 |
| 3831 | 98.75 | 99.35 |
| 3830 | 98.74 | 99.35 |
| 3829 | 98.74 | 99.35 |
| 3828 | 98.74 | 99.34 |
| 3827 | 98.75 | 99.34 |
| 3826 | 98.77 | 99.34 |
| 3825 | 98.8 | 99.34 |
| 3824 | 98.8 | 99.34 |
| 3823 | 98.79 | 99.35 |
| 3822 | 98.77 | 99.35 |
| 3821 | 98.77 | 99.34 |
| 3820 | 98.79 | 99.33 |
| 3819 | 98.81 | 99.31 |
| 3818 | 98.81 | 99.31 |
| 3817 | 98.79 | 99.32 |
| 3816 | 98.79 | 99.34 |
| 3815 | 98.78 | 99.36 |
| 3814 | 98.77 | 99.38 |
| 3813 | 98.75 | 99.38 |
| 3812 | 98.74 | 99.39 |
| 3811 | 98.74 | 99.39 |
| 3810 | 98.76 | 99.39 |
| 3809 | 98.78 | 99.38 |
| 3808 | 98.79 | 99.38 |
| 3807 | 98.8 | 99.37 |
| 3806 | 98.8 | 99.36 |
| 3805 | 98.8 | 99.35 |
| 3804 | 98.79 | 99.35 |
| 3803 | 98.78 | 99.35 |
| 3802 | 98.78 | 99.35 |
| 3801 | 98.79 | 99.36 |

Transmittance (%)

| Wavenumber (cm-1) | ELP-RGD | ELP RGD azide |
| --- | --- | --- |
| 4000 | 98.99 | 98.42 |
| 3999 | 98.99 | 98.43 |
| 3998 | 98.99 | 98.43 |
| 3997 | 98.98 | 98.44 |
| 3996 | 98.97 | 98.44 |
| 3995 | 98.96 | 98.43 |
| 3994 | 98.96 | 98.43 |
| 3993 | 98.95 | 98.43 |
| 3992 | 98.95 | 98.44 |
| 3991 | 98.95 | 98.45 |
| 3990 | 98.96 | 98.46 |
| 3989 | 98.97 | 98.46 |
| 3988 | 98.97 | 98.44 |
| 3987 | 98.98 | 98.43 |
| 3986 | 99 | 98.41 |
| 3985 | 99.01 | 98.4 |
| 3984 | 99.02 | 98.4 |
| 3983 | 99.02 | 98.39 |
| 3982 | 99.01 | 98.39 |
| 3981 | 98.99 | 98.39 |
| 3980 | 98.98 | 98.4 |
| 3979 | 98.98 | 98.41 |
| 3978 | 98.99 | 98.42 |
| 3977 | 99 | 98.42 |
| 3976 | 99.02 | 98.43 |
| 3975 | 99.02 | 98.43 |
| 3974 | 99.03 | 98.43 |
| 3973 | 99.02 | 98.43 |
| 3972 | 99.02 | 98.43 |
| 3971 | 99.01 | 98.42 |
| 3970 | 99 | 98.41 |
| 3969 | 99 | 98.4 |
| 3968 | 98.99 | 98.39 |
| 3967 | 98.99 | 98.39 |
| 3966 | 98.99 | 98.39 |
| 3965 | 98.99 | 98.41 |
| 3964 | 98.99 | 98.44 |
| 3963 | 98.99 | 98.45 |
| 3962 | 98.99 | 98.46 |
| 3961 | 99 | 98.45 |
| 3960 | 99 | 98.43 |
| 3959 | 99 | 98.42 |
| 3958 | 98.99 | 98.41 |
| 3957 | 98.98 | 98.42 |
| 3956 | 98.98 | 98.43 |
| 3955 | 98.98 | 98.44 |
| 3954 | 98.98 | 98.45 |
| 3953 | 98.99 | 98.44 |
| 3952 | 98.99 | 98.42 |
| 3951 | 98.99 | 98.4 |
| 3950 | 98.99 | 98.38 |
| 3949 | 98.99 | 98.38 |
| 3948 | 99 | 98.38 |
| 3947 | 99 | 98.39 |
| 3946 | 99.01 | 98.41 |
| 3945 | 99.02 | 98.42 |
| 3944 | 99.03 | 98.42 |
| 3943 | 99.03 | 98.42 |
| 3942 | 99.03 | 98.4 |
| 3941 | 99.03 | 98.39 |
| 3940 | 99.03 | 98.38 |
| 3939 | 99.03 | 98.38 |
| 3938 | 99.04 | 98.39 |
| 3937 | 99.04 | 98.41 |
| 3936 | 99.05 | 98.41 |
| 3935 | 99.04 | 98.41 |
| 3934 | 99.04 | 98.4 |
| 3933 | 99.03 | 98.39 |
| 3932 | 99.03 | 98.38 |
| 3931 | 99.02 | 98.38 |
| 3930 | 99.02 | 98.39 |
| 3929 | 99.03 | 98.4 |
| 3928 | 99.03 | 98.42 |
| 3927 | 99.04 | 98.43 |
| 3926 | 99.04 | 98.43 |
| 3925 | 99.05 | 98.42 |
| 3924 | 99.05 | 98.41 |
| 3923 | 99.04 | 98.4 |
| 3922 | 99.04 | 98.4 |
| 3921 | 99.03 | 98.39 |
| 3920 | 99.03 | 98.39 |
| 3919 | 99.02 | 98.37 |
| 3918 | 99.01 | 98.36 |
| 3917 | 99.01 | 98.35 |
| 3916 | 99.02 | 98.35 |
| 3915 | 99.02 | 98.37 |
| 3914 | 99.03 | 98.39 |
| 3913 | 99.04 | 98.41 |
| 3912 | 99.05 | 98.42 |
| 3911 | 99.06 | 98.43 |
| 3910 | 99.07 | 98.41 |
| 3909 | 99.07 | 98.39 |
| 3908 | 99.06 | 98.37 |
| 3907 | 99.04 | 98.35 |
| 3906 | 99.02 | 98.34 |
| 3905 | 99.01 | 98.34 |
| 3904 | 99.03 | 98.34 |
| 3903 | 99.04 | 98.35 |
| 3902 | 99.06 | 98.37 |
| 3901 | 99.07 | 98.39 |
| 3900 | 99.06 | 98.4 |
| 3899 | 99.05 | 98.42 |
| 3898 | 99.04 | 98.42 |
| 3897 | 99.03 | 98.42 |
| 3896 | 99.03 | 98.42 |
| 3895 | 99.03 | 98.41 |
| 3894 | 99.03 | 98.41 |
| 3893 | 99.02 | 98.42 |
| 3892 | 99.02 | 98.42 |
| 3891 | 99.02 | 98.42 |
| 3890 | 99.02 | 98.41 |
| 3889 | 99.02 | 98.41 |
| 3888 | 99.03 | 98.41 |
| 3887 | 99.04 | 98.4 |
| 3886 | 99.04 | 98.39 |
| 3885 | 99.04 | 98.38 |
| 3884 | 99.03 | 98.38 |
| 3883 | 99.02 | 98.38 |
| 3882 | 99 | 98.39 |
| 3881 | 98.98 | 98.4 |
| 3880 | 98.97 | 98.41 |
| 3879 | 98.97 | 98.42 |
| 3878 | 98.98 | 98.42 |
| 3877 | 99 | 98.41 |
| 3876 | 99.02 | 98.4 |
| 3875 | 99.04 | 98.4 |
| 3874 | 99.06 | 98.4 |
| 3873 | 99.07 | 98.41 |
| 3872 | 99.07 | 98.41 |
| 3871 | 99.07 | 98.4 |
| 3870 | 99.07 | 98.39 |
| 3869 | 99.06 | 98.38 |
| 3868 | 99.05 | 98.38 |
| 3867 | 99.04 | 98.38 |
| 3866 | 99.04 | 98.37 |
| 3865 | 99.04 | 98.36 |
| 3864 | 99.05 | 98.36 |
| 3863 | 99.05 | 98.35 |
| 3862 | 99.05 | 98.36 |
| 3861 | 99.05 | 98.37 |
| 3860 | 99.05 | 98.38 |
| 3859 | 99.06 | 98.39 |
| 3858 | 99.06 | 98.39 |
| 3857 | 99.07 | 98.39 |
| 3856 | 99.07 | 98.38 |
| 3855 | 99.06 | 98.38 |
| 3854 | 99.06 | 98.37 |
| 3853 | 99.06 | 98.35 |
| 3852 | 99.05 | 98.32 |
| 3851 | 99.04 | 98.3 |
| 3850 | 99.03 | 98.31 |
| 3849 | 99.04 | 98.34 |
| 3848 | 99.05 | 98.39 |
| 3847 | 99.07 | 98.43 |
| 3846 | 99.09 | 98.45 |
| 3845 | 99.11 | 98.45 |
| 3844 | 99.11 | 98.44 |
| 3843 | 99.1 | 98.43 |
| 3842 | 99.09 | 98.42 |
| 3841 | 99.08 | 98.42 |
| 3840 | 99.08 | 98.41 |
| 3839 | 99.08 | 98.39 |
| 3838 | 99.07 | 98.38 |
| 3837 | 99.06 | 98.39 |
| 3836 | 99.06 | 98.4 |
| 3835 | 99.06 | 98.41 |
| 3834 | 99.08 | 98.42 |
| 3833 | 99.09 | 98.43 |
| 3832 | 99.09 | 98.44 |
| 3831 | 99.07 | 98.45 |
| 3830 | 99.06 | 98.45 |
| 3829 | 99.04 | 98.44 |
| 3828 | 99.03 | 98.44 |
| 3827 | 99.03 | 98.43 |
| 3826 | 99.04 | 98.41 |
| 3825 | 99.04 | 98.4 |
| 3824 | 99.05 | 98.39 |
| 3823 | 99.06 | 98.39 |
| 3822 | 99.07 | 98.39 |
| 3821 | 99.08 | 98.41 |
| 3820 | 99.08 | 98.42 |
| 3819 | 99.09 | 98.42 |
| 3818 | 99.1 | 98.42 |
| 3817 | 99.11 | 98.41 |
| 3816 | 99.11 | 98.42 |
| 3815 | 99.11 | 98.44 |
| 3814 | 99.1 | 98.45 |
| 3813 | 99.09 | 98.44 |
| 3812 | 99.09 | 98.43 |
| 3811 | 99.09 | 98.43 |
| 3810 | 99.09 | 98.44 |
| 3809 | 99.09 | 98.45 |
| 3808 | 99.1 | 98.46 |
| 3807 | 99.11 | 98.46 |
| 3806 | 99.11 | 98.46 |
| 3805 | 99.1 | 98.44 |
| 3804 | 99.1 | 98.43 |
| 3803 | 99.09 | 98.41 |
| 3802 | 99.08 | 98.41 |
| 3801 | 99.07 | 98.41 |

Transmittance (%)

| Wavenumber (cm-1) | ELP-RDG | ELP-RDG azide |
| --- | --- | --- |
| 4000 | 98.89 | 98.69 |
| 3999 | 98.9 | 98.71 |
| 3998 | 98.9 | 98.72 |
| 3997 | 98.91 | 98.73 |
| 3996 | 98.91 | 98.73 |
| 3995 | 98.91 | 98.72 |
| 3994 | 98.9 | 98.71 |
| 3993 | 98.89 | 98.71 |
| 3992 | 98.88 | 98.71 |
| 3991 | 98.88 | 98.71 |
| 3990 | 98.89 | 98.72 |
| 3989 | 98.9 | 98.73 |
| 3988 | 98.9 | 98.74 |
| 3987 | 98.9 | 98.73 |
| 3986 | 98.9 | 98.73 |
| 3985 | 98.9 | 98.72 |
| 3984 | 98.9 | 98.71 |
| 3983 | 98.9 | 98.71 |
| 3982 | 98.9 | 98.71 |
| 3981 | 98.91 | 98.71 |
| 3980 | 98.91 | 98.72 |
| 3979 | 98.92 | 98.73 |
| 3978 | 98.92 | 98.73 |
| 3977 | 98.92 | 98.72 |
| 3976 | 98.92 | 98.72 |
| 3975 | 98.92 | 98.72 |
| 3974 | 98.92 | 98.73 |
| 3973 | 98.92 | 98.74 |
| 3972 | 98.91 | 98.74 |
| 3971 | 98.9 | 98.74 |
| 3970 | 98.9 | 98.73 |
| 3969 | 98.89 | 98.73 |
| 3968 | 98.9 | 98.72 |
| 3967 | 98.9 | 98.72 |
| 3966 | 98.91 | 98.72 |
| 3965 | 98.92 | 98.72 |
| 3964 | 98.92 | 98.72 |
| 3963 | 98.92 | 98.72 |
| 3962 | 98.92 | 98.72 |
| 3961 | 98.91 | 98.72 |
| 3960 | 98.9 | 98.73 |
| 3959 | 98.89 | 98.73 |
| 3958 | 98.88 | 98.74 |
| 3957 | 98.87 | 98.74 |
| 3956 | 98.87 | 98.74 |
| 3955 | 98.88 | 98.73 |
| 3954 | 98.89 | 98.72 |
| 3953 | 98.89 | 98.72 |
| 3952 | 98.9 | 98.71 |
| 3951 | 98.9 | 98.7 |
| 3950 | 98.91 | 98.69 |
| 3949 | 98.91 | 98.67 |
| 3948 | 98.92 | 98.66 |
| 3947 | 98.93 | 98.66 |
| 3946 | 98.94 | 98.67 |
| 3945 | 98.94 | 98.68 |
| 3944 | 98.95 | 98.69 |
| 3943 | 98.96 | 98.69 |
| 3942 | 98.96 | 98.69 |
| 3941 | 98.96 | 98.69 |
| 3940 | 98.96 | 98.68 |
| 3939 | 98.96 | 98.68 |
| 3938 | 98.95 | 98.69 |
| 3937 | 98.94 | 98.69 |
| 3936 | 98.93 | 98.69 |
| 3935 | 98.93 | 98.68 |
| 3934 | 98.93 | 98.68 |
| 3933 | 98.95 | 98.69 |
| 3932 | 98.96 | 98.7 |
| 3931 | 98.97 | 98.71 |
| 3930 | 98.98 | 98.72 |
| 3929 | 98.98 | 98.72 |
| 3928 | 98.97 | 98.72 |
| 3927 | 98.95 | 98.72 |
| 3926 | 98.94 | 98.72 |
| 3925 | 98.92 | 98.72 |
| 3924 | 98.91 | 98.72 |
| 3923 | 98.92 | 98.72 |
| 3922 | 98.94 | 98.71 |
| 3921 | 98.96 | 98.7 |
| 3920 | 98.97 | 98.7 |
| 3919 | 98.98 | 98.7 |
| 3918 | 98.99 | 98.7 |
| 3917 | 98.99 | 98.71 |
| 3916 | 98.99 | 98.72 |
| 3915 | 98.98 | 98.72 |
| 3914 | 98.98 | 98.73 |
| 3913 | 98.97 | 98.73 |
| 3912 | 98.97 | 98.72 |
| 3911 | 98.98 | 98.71 |
| 3910 | 98.98 | 98.69 |
| 3909 | 98.98 | 98.68 |
| 3908 | 98.98 | 98.68 |
| 3907 | 98.97 | 98.68 |
| 3906 | 98.96 | 98.69 |
| 3905 | 98.97 | 98.7 |
| 3904 | 98.97 | 98.7 |
| 3903 | 98.97 | 98.69 |
| 3902 | 98.97 | 98.69 |
| 3901 | 98.96 | 98.68 |
| 3900 | 98.96 | 98.69 |
| 3899 | 98.96 | 98.7 |
| 3898 | 98.97 | 98.7 |
| 3897 | 98.96 | 98.7 |
| 3896 | 98.96 | 98.69 |
| 3895 | 98.95 | 98.68 |
| 3894 | 98.96 | 98.68 |
| 3893 | 98.96 | 98.67 |
| 3892 | 98.97 | 98.68 |
| 3891 | 98.97 | 98.68 |
| 3890 | 98.96 | 98.7 |
| 3889 | 98.96 | 98.7 |
| 3888 | 98.97 | 98.7 |
| 3887 | 98.98 | 98.71 |
| 3886 | 98.99 | 98.71 |
| 3885 | 98.99 | 98.71 |
| 3884 | 98.98 | 98.7 |
| 3883 | 98.96 | 98.7 |
| 3882 | 98.94 | 98.69 |
| 3881 | 98.93 | 98.7 |
| 3880 | 98.93 | 98.71 |
| 3879 | 98.92 | 98.72 |
| 3878 | 98.92 | 98.71 |
| 3877 | 98.92 | 98.7 |
| 3876 | 98.93 | 98.68 |
| 3875 | 98.94 | 98.67 |
| 3874 | 98.95 | 98.68 |
| 3873 | 98.96 | 98.67 |
| 3872 | 98.96 | 98.66 |
| 3871 | 98.96 | 98.65 |
| 3870 | 98.97 | 98.65 |
| 3869 | 98.97 | 98.65 |
| 3868 | 98.97 | 98.65 |
| 3867 | 98.97 | 98.64 |
| 3866 | 98.98 | 98.63 |
| 3865 | 98.99 | 98.64 |
| 3864 | 98.99 | 98.65 |
| 3863 | 98.99 | 98.65 |
| 3862 | 98.98 | 98.66 |
| 3861 | 98.98 | 98.67 |
| 3860 | 98.98 | 98.69 |
| 3859 | 98.99 | 98.71 |
| 3858 | 98.99 | 98.72 |
| 3857 | 98.99 | 98.73 |
| 3856 | 98.98 | 98.73 |
| 3855 | 98.98 | 98.73 |
| 3854 | 98.98 | 98.73 |
| 3853 | 98.98 | 98.72 |
| 3852 | 98.99 | 98.71 |
| 3851 | 98.99 | 98.69 |
| 3850 | 99 | 98.69 |
| 3849 | 99 | 98.7 |
| 3848 | 99 | 98.72 |
| 3847 | 99.01 | 98.73 |
| 3846 | 99.02 | 98.73 |
| 3845 | 99.02 | 98.72 |
| 3844 | 99.01 | 98.71 |
| 3843 | 98.99 | 98.71 |
| 3842 | 98.98 | 98.72 |
| 3841 | 98.97 | 98.72 |
| 3840 | 98.96 | 98.71 |
| 3839 | 98.96 | 98.7 |
| 3838 | 98.96 | 98.69 |
| 3837 | 98.96 | 98.69 |
| 3836 | 98.97 | 98.7 |
| 3835 | 98.99 | 98.71 |
| 3834 | 99.01 | 98.73 |
| 3833 | 99.02 | 98.74 |
| 3832 | 99.02 | 98.76 |
| 3831 | 99.01 | 98.78 |
| 3830 | 99 | 98.78 |
| 3829 | 98.99 | 98.77 |
| 3828 | 98.98 | 98.76 |
| 3827 | 98.98 | 98.75 |
| 3826 | 98.98 | 98.74 |
| 3825 | 98.98 | 98.74 |
| 3824 | 98.98 | 98.74 |
| 3823 | 98.98 | 98.74 |
| 3822 | 98.99 | 98.75 |
| 3821 | 98.99 | 98.75 |
| 3820 | 99 | 98.74 |
| 3819 | 99 | 98.73 |
| 3818 | 99 | 98.72 |
| 3817 | 99 | 98.72 |
| 3816 | 98.99 | 98.74 |
| 3815 | 98.98 | 98.76 |
| 3814 | 98.97 | 98.76 |
| 3813 | 98.96 | 98.74 |
| 3812 | 98.95 | 98.72 |
| 3811 | 98.95 | 98.71 |
| 3810 | 98.96 | 98.71 |
| 3809 | 98.97 | 98.73 |
| 3808 | 98.98 | 98.76 |
| 3807 | 98.99 | 98.78 |
| 3806 | 98.98 | 98.78 |
| 3805 | 98.99 | 98.76 |
| 3804 | 99 | 98.73 |
| 3803 | 99.02 | 98.71 |
| 3802 | 99.03 | 98.69 |
| 3801 | 99.02 | 98.7 |

Transmittance (%)

| Wavenumber (cm-1) | ELP-RGD | ELP-RGD azide (ave 3.78) | ELP-RGD azide (ave 12) |
| --- | --- | --- | --- |
| 4000 | 98.99 | 97.62 | 98.2 |
| 3999 | 98.99 | 97.62 | 98.21 |
| 3998 | 98.99 | 97.61 | 98.21 |
| 3997 | 98.98 | 97.61 | 98.21 |
| 3996 | 98.97 | 97.61 | 98.2 |
| 3995 | 98.96 | 97.61 | 98.19 |
| 3994 | 98.96 | 97.6 | 98.18 |
| 3993 | 98.95 | 97.59 | 98.17 |
| 3992 | 98.95 | 97.58 | 98.16 |
| 3991 | 98.95 | 97.58 | 98.15 |
| 3990 | 98.96 | 97.58 | 98.15 |
| 3989 | 98.97 | 97.59 | 98.15 |
| 3988 | 98.97 | 97.6 | 98.15 |
| 3987 | 98.98 | 97.6 | 98.15 |
| 3986 | 99 | 97.6 | 98.16 |
| 3985 | 99.01 | 97.6 | 98.16 |
| 3984 | 99.02 | 97.61 | 98.16 |
| 3983 | 99.02 | 97.61 | 98.16 |
| 3982 | 99.01 | 97.6 | 98.16 |
| 3981 | 98.99 | 97.59 | 98.17 |
| 3980 | 98.98 | 97.59 | 98.18 |
| 3979 | 98.98 | 97.59 | 98.19 |
| 3978 | 98.99 | 97.6 | 98.19 |
| 3977 | 99 | 97.61 | 98.19 |
| 3976 | 99.02 | 97.63 | 98.2 |
| 3975 | 99.02 | 97.63 | 98.2 |
| 3974 | 99.03 | 97.64 | 98.21 |
| 3973 | 99.02 | 97.63 | 98.21 |
| 3972 | 99.02 | 97.62 | 98.2 |
| 3971 | 99.01 | 97.61 | 98.19 |
| 3970 | 99 | 97.6 | 98.18 |
| 3969 | 99 | 97.6 | 98.17 |
| 3968 | 98.99 | 97.59 | 98.17 |
| 3967 | 98.99 | 97.59 | 98.16 |
| 3966 | 98.99 | 97.6 | 98.17 |
| 3965 | 98.99 | 97.62 | 98.17 |
| 3964 | 98.99 | 97.64 | 98.17 |
| 3963 | 98.99 | 97.65 | 98.18 |
| 3962 | 98.99 | 97.66 | 98.18 |
| 3961 | 99 | 97.66 | 98.2 |
| 3960 | 99 | 97.66 | 98.2 |
| 3959 | 99 | 97.65 | 98.21 |
| 3958 | 98.99 | 97.65 | 98.2 |
| 3957 | 98.98 | 97.64 | 98.2 |
| 3956 | 98.98 | 97.64 | 98.2 |
| 3955 | 98.98 | 97.63 | 98.21 |
| 3954 | 98.98 | 97.63 | 98.22 |
| 3953 | 98.99 | 97.63 | 98.23 |
| 3952 | 98.99 | 97.63 | 98.22 |
| 3951 | 98.99 | 97.64 | 98.2 |
| 3950 | 98.99 | 97.64 | 98.19 |
| 3949 | 98.99 | 97.64 | 98.18 |
| 3948 | 99 | 97.64 | 98.18 |
| 3947 | 99 | 97.64 | 98.18 |
| 3946 | 99.01 | 97.65 | 98.19 |
| 3945 | 99.02 | 97.65 | 98.21 |
| 3944 | 99.03 | 97.66 | 98.22 |
| 3943 | 99.03 | 97.66 | 98.23 |
| 3942 | 99.03 | 97.65 | 98.24 |
| 3941 | 99.03 | 97.65 | 98.24 |
| 3940 | 99.03 | 97.65 | 98.23 |
| 3939 | 99.03 | 97.66 | 98.22 |
| 3938 | 99.04 | 97.66 | 98.22 |
| 3937 | 99.04 | 97.67 | 98.22 |
| 3936 | 99.05 | 97.67 | 98.23 |
| 3935 | 99.04 | 97.67 | 98.25 |
| 3934 | 99.04 | 97.68 | 98.25 |
| 3933 | 99.03 | 97.68 | 98.25 |
| 3932 | 99.03 | 97.68 | 98.24 |
| 3931 | 99.02 | 97.68 | 98.22 |
| 3930 | 99.02 | 97.67 | 98.2 |
| 3929 | 99.03 | 97.66 | 98.2 |
| 3928 | 99.03 | 97.67 | 98.2 |
| 3927 | 99.04 | 97.69 | 98.21 |
| 3926 | 99.04 | 97.71 | 98.22 |
| 3925 | 99.05 | 97.71 | 98.24 |
| 3924 | 99.05 | 97.71 | 98.25 |
| 3923 | 99.04 | 97.71 | 98.26 |
| 3922 | 99.04 | 97.71 | 98.26 |
| 3921 | 99.03 | 97.71 | 98.26 |
| 3920 | 99.03 | 97.71 | 98.25 |
| 3919 | 99.02 | 97.71 | 98.25 |
| 3918 | 99.01 | 97.71 | 98.26 |
| 3917 | 99.01 | 97.71 | 98.28 |
| 3916 | 99.02 | 97.72 | 98.28 |
| 3915 | 99.02 | 97.72 | 98.27 |
| 3914 | 99.03 | 97.71 | 98.26 |
| 3913 | 99.04 | 97.69 | 98.25 |
| 3912 | 99.05 | 97.68 | 98.24 |
| 3911 | 99.06 | 97.68 | 98.24 |
| 3910 | 99.07 | 97.69 | 98.25 |
| 3909 | 99.07 | 97.69 | 98.26 |
| 3908 | 99.06 | 97.7 | 98.28 |
| 3907 | 99.04 | 97.7 | 98.28 |
| 3906 | 99.02 | 97.69 | 98.27 |
| 3905 | 99.01 | 97.69 | 98.27 |
| 3904 | 99.03 | 97.69 | 98.27 |
| 3903 | 99.04 | 97.7 | 98.29 |
| 3902 | 99.06 | 97.72 | 98.31 |
| 3901 | 99.07 | 97.73 | 98.32 |
| 3900 | 99.06 | 97.74 | 98.31 |
| 3899 | 99.05 | 97.75 | 98.3 |
| 3898 | 99.04 | 97.75 | 98.28 |
| 3897 | 99.03 | 97.75 | 98.26 |
| 3896 | 99.03 | 97.75 | 98.25 |
| 3895 | 99.03 | 97.75 | 98.25 |
| 3894 | 99.03 | 97.75 | 98.26 |
| 3893 | 99.02 | 97.74 | 98.28 |
| 3892 | 99.02 | 97.74 | 98.29 |
| 3891 | 99.02 | 97.74 | 98.29 |
| 3890 | 99.02 | 97.74 | 98.3 |
| 3889 | 99.02 | 97.74 | 98.31 |
| 3888 | 99.03 | 97.75 | 98.32 |
| 3887 | 99.04 | 97.75 | 98.33 |
| 3886 | 99.04 | 97.76 | 98.33 |
| 3885 | 99.04 | 97.76 | 98.32 |
| 3884 | 99.03 | 97.75 | 98.32 |
| 3883 | 99.02 | 97.73 | 98.32 |
| 3882 | 99 | 97.71 | 98.3 |
| 3881 | 98.98 | 97.71 | 98.29 |
| 3880 | 98.97 | 97.71 | 98.28 |
| 3879 | 98.97 | 97.72 | 98.27 |
| 3878 | 98.98 | 97.73 | 98.28 |
| 3877 | 99 | 97.72 | 98.28 |
| 3876 | 99.02 | 97.71 | 98.27 |
| 3875 | 99.04 | 97.71 | 98.27 |
| 3874 | 99.06 | 97.72 | 98.28 |
| 3873 | 99.07 | 97.74 | 98.3 |
| 3872 | 99.07 | 97.76 | 98.33 |
| 3871 | 99.07 | 97.77 | 98.34 |
| 3870 | 99.07 | 97.78 | 98.35 |
| 3869 | 99.06 | 97.78 | 98.36 |
| 3868 | 99.05 | 97.77 | 98.36 |
| 3867 | 99.04 | 97.76 | 98.35 |
| 3866 | 99.04 | 97.76 | 98.33 |
| 3865 | 99.04 | 97.76 | 98.32 |
| 3864 | 99.05 | 97.78 | 98.32 |
| 3863 | 99.05 | 97.79 | 98.33 |
| 3862 | 99.05 | 97.79 | 98.33 |
| 3861 | 99.05 | 97.79 | 98.32 |
| 3860 | 99.05 | 97.78 | 98.32 |
| 3859 | 99.06 | 97.78 | 98.32 |
| 3858 | 99.06 | 97.79 | 98.34 |
| 3857 | 99.07 | 97.8 | 98.35 |
| 3856 | 99.07 | 97.8 | 98.36 |
| 3855 | 99.06 | 97.79 | 98.35 |
| 3854 | 99.06 | 97.78 | 98.35 |
| 3853 | 99.06 | 97.8 | 98.35 |
| 3852 | 99.05 | 97.81 | 98.36 |
| 3851 | 99.04 | 97.81 | 98.35 |
| 3850 | 99.03 | 97.8 | 98.34 |
| 3849 | 99.04 | 97.79 | 98.33 |
| 3848 | 99.05 | 97.79 | 98.33 |
| 3847 | 99.07 | 97.78 | 98.33 |
| 3846 | 99.09 | 97.78 | 98.33 |
| 3845 | 99.11 | 97.78 | 98.33 |
| 3844 | 99.11 | 97.79 | 98.33 |
| 3843 | 99.1 | 97.8 | 98.33 |
| 3842 | 99.09 | 97.81 | 98.34 |
| 3841 | 99.08 | 97.82 | 98.35 |
| 3840 | 99.08 | 97.83 | 98.35 |
| 3839 | 99.08 | 97.83 | 98.35 |
| 3838 | 99.07 | 97.82 | 98.35 |
| 3837 | 99.06 | 97.82 | 98.36 |
| 3836 | 99.06 | 97.81 | 98.37 |
| 3835 | 99.06 | 97.8 | 98.38 |
| 3834 | 99.08 | 97.8 | 98.37 |
| 3833 | 99.09 | 97.79 | 98.36 |
| 3832 | 99.09 | 97.79 | 98.34 |
| 3831 | 99.07 | 97.79 | 98.33 |
| 3830 | 99.06 | 97.79 | 98.32 |
| 3829 | 99.04 | 97.79 | 98.33 |
| 3828 | 99.03 | 97.79 | 98.33 |
| 3827 | 99.03 | 97.79 | 98.33 |
| 3826 | 99.04 | 97.79 | 98.34 |
| 3825 | 99.04 | 97.81 | 98.36 |
| 3824 | 99.05 | 97.82 | 98.38 |
| 3823 | 99.06 | 97.82 | 98.39 |
| 3822 | 99.07 | 97.83 | 98.39 |
| 3821 | 99.08 | 97.84 | 98.39 |
| 3820 | 99.08 | 97.85 | 98.4 |
| 3819 | 99.09 | 97.85 | 98.4 |
| 3818 | 99.1 | 97.85 | 98.39 |
| 3817 | 99.11 | 97.84 | 98.37 |
| 3816 | 99.11 | 97.84 | 98.35 |
| 3815 | 99.11 | 97.84 | 98.35 |
| 3814 | 99.1 | 97.83 | 98.34 |
| 3813 | 99.09 | 97.82 | 98.34 |
| 3812 | 99.09 | 97.81 | 98.33 |
| 3811 | 99.09 | 97.81 | 98.33 |
| 3810 | 99.09 | 97.81 | 98.35 |
| 3809 | 99.09 | 97.81 | 98.38 |
| 3808 | 99.1 | 97.81 | 98.4 |
| 3807 | 99.11 | 97.82 | 98.4 |
| 3806 | 99.11 | 97.83 | 98.4 |
| 3805 | 99.1 | 97.84 | 98.4 |
| 3804 | 99.1 | 97.85 | 98.41 |
| 3803 | 99.09 | 97.85 | 98.4 |
| 3802 | 99.08 | 97.84 | 98.39 |
| 3801 | 99.07 | 97.83 | 98.38 |

**Supplemental Figure 1**: FTIR of ELP pre- and post-azide functionalization. The appearance of a peak at 2100 cm^-1^ for (A) uPA degradable sequence ELP-u1, (B) integrin binding sequence ELP-RGD, and (C) non-bioactive sequence ELP-RDG indicated the presence of azide bonds within the protein structure. (D) The azide peak intensity increased with higher azide functionalization as indicated by FTIR of ELP-RGD sequence pre-modification and protein batches modified with an average of 3.78 and 12 azides. The average azide group addition was controlled by stoichiometrically limiting the number of azides present during the diazotransfer reaction.

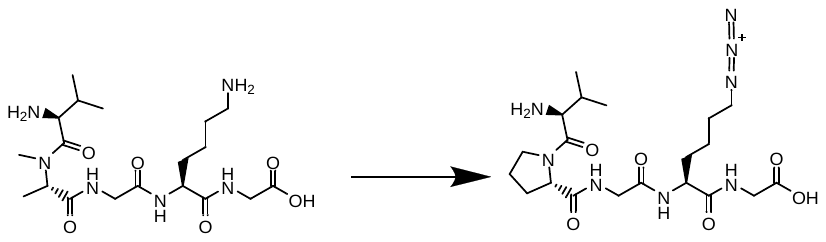

A

ELP (azide functionalized)

ELP (-VPG**K**G-)

26 Da Mw increase per azide

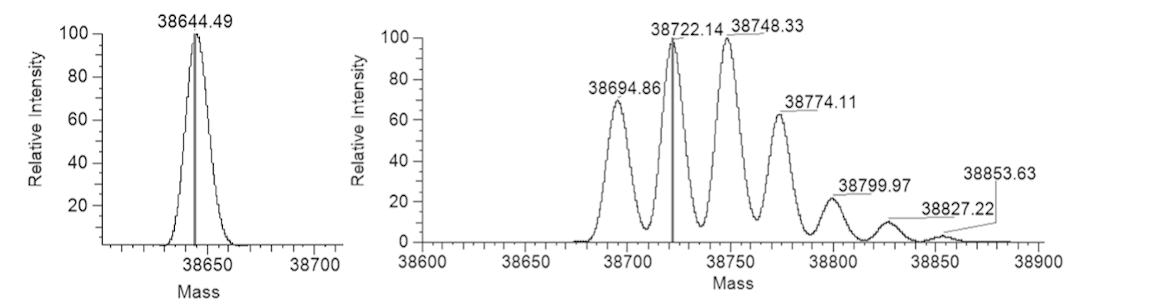

ELP-RGD azide

ELP-RGD

C

ELP-u1 azide

ELP-u1

B

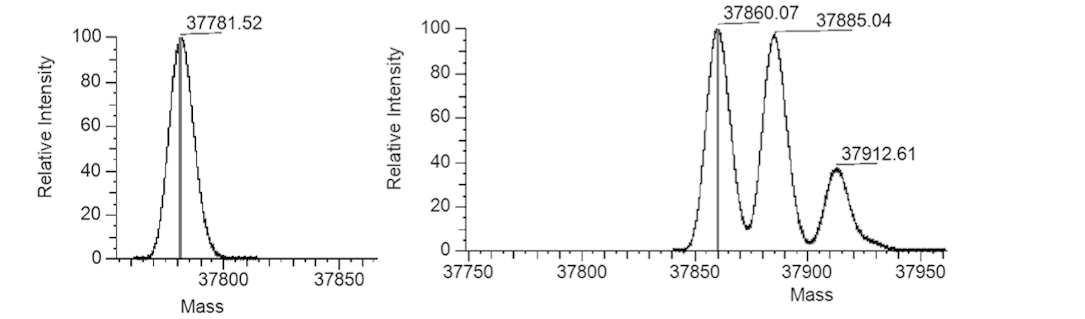

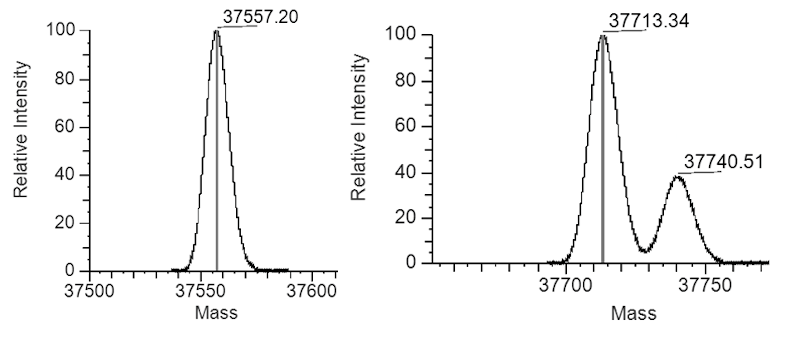

ELP-RDG azide

D

ELP-RDG

**Supplemental Figure 2**: High resolution electrospray ionization-mass spectrometry (ESI-MS) of ELP pre- and post-azide functionalization. (A) Each lysine residue modification with azides increases protein mass by 26 Da. Pre-azide modification ELP sequence appears as one peak, indicating monodisperse protein molecular weight. Post-azide modification ELP appears as multiple peaks with 26 Da differences, indicating that the diazo transfer reaction created ELPs with a distribution of attached azides: (B) uPA degradable sequence ELP-u1 pre- and post-azide functionalization, (C) cell-adhesive sequence ELP-RGD pre- and post-azide functionalization, and (D) non-bioactive sequence ELP-RDG pre- and post-azide functionalization. The relative peak intensities were quantified to determine the molecular average of azides added to ELP-u1, -RGD and –RDG, which were 6.28, 3.78, and 3.74, respectively.

**Supplemental Figure 3**: THPC used to immobilize ELP to poly(acrylamide) gel for zymography. (A) Poly(acrylamide) crosslinking chemistry creates a gel with primary amines in the structure. (B) THPC has four primary amine reactive sites that allow for ELP to be immobilized onto poly(acrylamide) gel network. (C) Poly(acrylamide) gel dosed with ELP without THPC causes ELP to migrate down the wells during electrophoresis. (D) Poly(acrylamide gel dosed with immobilized ELP maintains uniform protein distribution after electrophoresis.

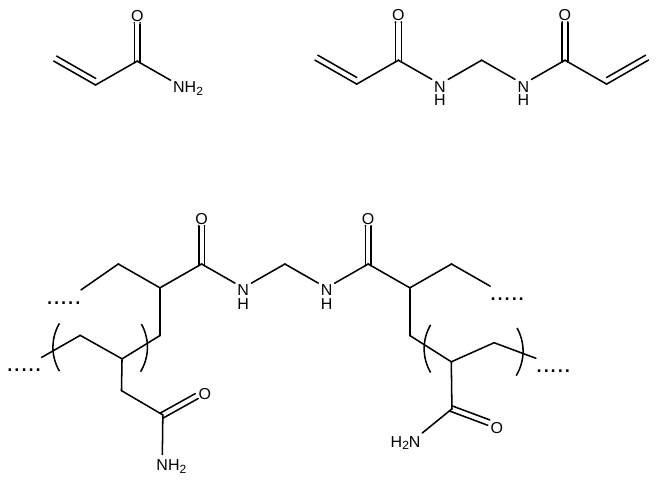

TEMED

APS

Acrylamide

Bis-acrylamide

ELP

immobilization site

A

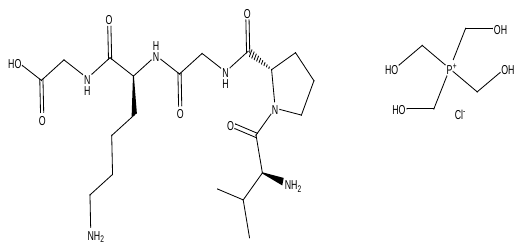

ELP (-VPG**K**G-)

THPC

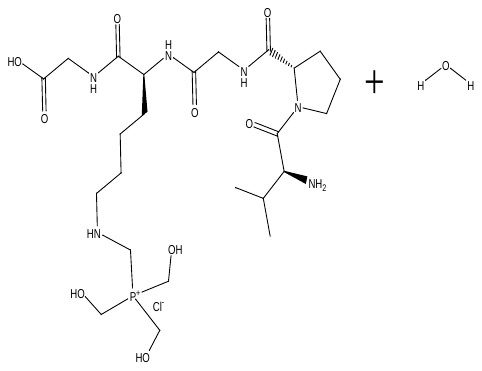

possible acrylamide gel binding site(s)

B

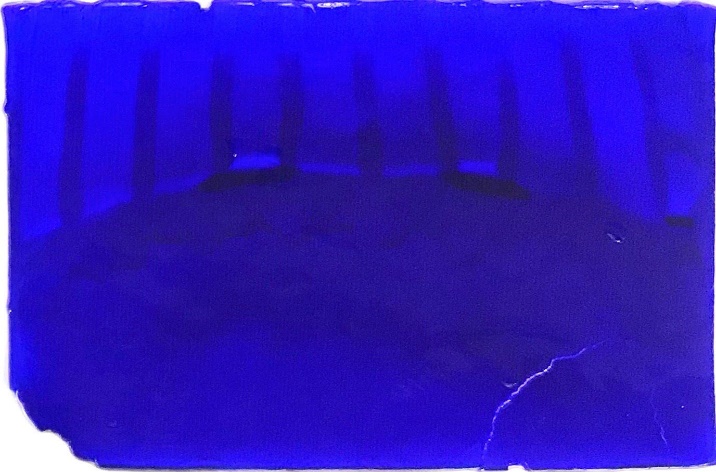

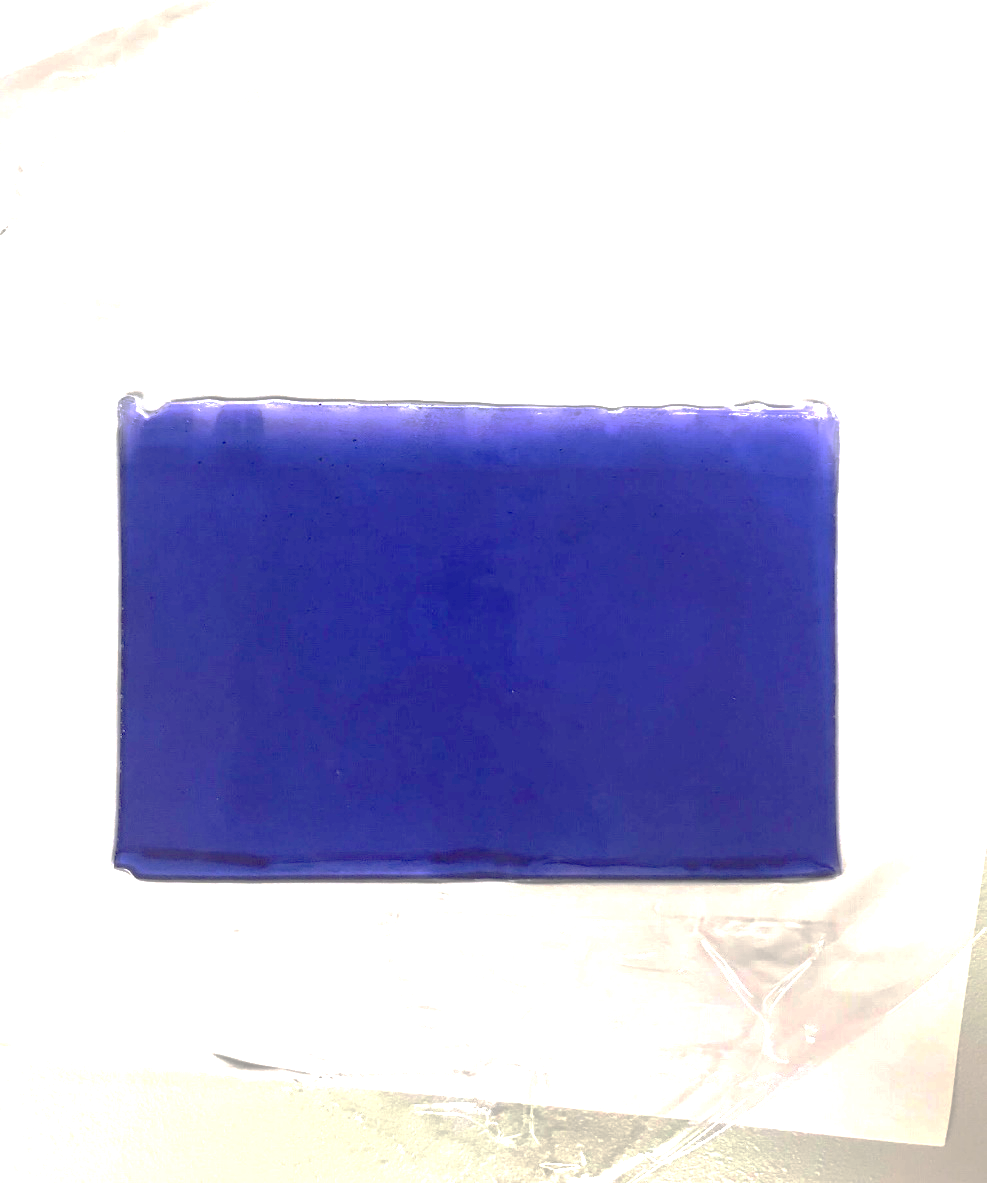

C

D

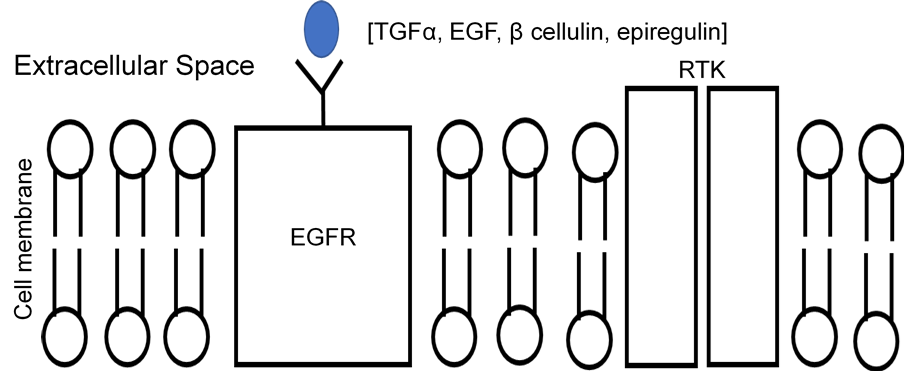

(Growth inhibition)

Cytosol

rapamycin

Nf1

U0126

SGK

mTORC1

Akt

PI3K

IRS-1

SO5

Grb2

Neurofibromin

p53

PTEN

Ras

Raf

MEK 1 & 2

ERK 1 & 2

pERK 1 & 2

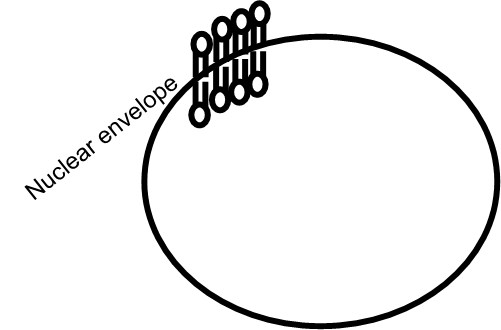

**Supplemental Figure 4**: OPC differentiation relevant pathways are impacted by the absence of Nf1 and p53. Disruption of ERK 1 & 2 pathway results in delayed OPC differentiation *in vivo*. Solid arrows indicate pathway promotion, solid bars indicate pathway inhibition, dashed lines indicate pathways not occurring in MADM OPCs. Red bars indicate drugs used to assess changes to OPC differentiation *in vivo* from the disruption of the ErK 1 & 2 and mTORC1 pathways.

**Supplemental Figure 4**: OPC differentiation relevant pathways are impacted by the absence of Nf1 and p53. Disruption of ERK 1 & 2 pathway results in delayed OPC differentiation *in vivo*. Solid arrows indicate pathway promotion, solid bars indicate pathway inhibition, dashed lines indicate pathways not occurring in MADM OPCs. Red bars indicate drugs used to assess changes to OPC differentiation *in vivo* from the disruption of the ErK 1 & 2 and mTORC1 pathways.
